# Aging-associated Decline in Vascular Smooth Muscle Cell Mechanosensation is Mediated by Piezo1 Channel

**DOI:** 10.1101/2023.04.27.538557

**Authors:** Ngoc Luu, Apratim Bajpai, Rui Li, Seojin Park, Mahad Noor, Xiao Ma, Weiqiang Chen

## Abstract

Aging of the vasculature is associated with detrimental changes in vascular smooth muscle cell (VSMC) mechanosensitivity to extrinsic forces in their surrounding microenvironment. However, how chronological aging alters VSMCs’ ability to sense and adapt to mechanical perturbations remains unexplored. Here, we show defective VSMC mechanosensation in aging measured with ultrasound tweezers-based micromechanical system, force instantaneous frequency spectrum and transcriptome analyses. The mechanobiological study reveals that aged VSMCs adapt a relatively inert solid-like state with altered actin cytoskeletal integrity, resulting in an impairment in their mechanosensitivity and dynamic mechanoresponse to mechanical perturbations. The aging-associated decline in mechanosensation behaviors is mediated by hyperactivity of Piezo1-dependent calcium signaling. Inhibition of Piezo1 alleviates vascular aging and partially restores the loss in dynamic contractile properties in aged cells. Altogether, our study reveals the novel signaling pathway underlying aging-associated aberrant mechanosensation in VSMC and identifies Piezo1 as a potential therapeutic mechanobiological target to alleviate vascular aging.

## Introduction

Aging is a major risk factor for cardiovascular diseases, such as atherosclerosis [1], aneurysms [2], atrial fibrillation [3], hypertension [4], and cardiac hypertrophy [5]. At both tissue and cellular levels, aging develops with detrimental changes in the structure and function of the vasculature, attributing to a notable decline in mechanical properties of vascular cells [6–8]. The aging of arterial wall impairs the natural ability to adapt to the rapidly changing mechanical stress (e.g., shear stress and cyclic stretch) exerted by blood flow due to the reducing mechanical responses of vascular cells in aorta. As the major motivators of aorta, vascular smooth muscle cells (VSMCs) are crucial in regulating the contractility of vessel walls and responding mechanical stress [9]. Thus, a healthy VSMC mechanosensation behavior (the capability of mechano-sensing and transduction of mechanical force into intracellular signals) enables the vessels to accommodate the rapidly changing mechanical perturbations from blood flows in arterial microenvironments. Stiffening of VSMC and dysfunctions in VSMC mechanosensation are often hallmarks of cardiovascular disease development [10–12]. Previous efforts of examining static mechanical properties of VSMC, such as stiffness and contractile force [13, 14], but not the dynamics mechanosensation activities, failed to reveal the aging-associated disease pathogenesis and progression mechanisms. Thus, understanding of the regulatory mechanisms of abnormal VSMC mechanosensation during aging and its role in aging and aging-associated cardiovascular diseases is critically needed yet currently limited.

Plenty of evidence have suggested that aging leads to changes in mechanical properties of VSMCs through a mechanobiological machinery. Cellular mechanosensation involves both mechanosensitive ion-gated channels or surface ligands to sense the extracellular mechanical cues, and actomyosin cytoskeleton (CSK) as a key mediator to generate intracellular mechanical forces and transmit such signals to the downstream mechano-responsive components [15–17]. VSMC mechanosensation depends on the swift opening of Piezo1, a tension-gated ion channel that allows the influx of calcium (Ca^2+^) ions upon activation to regulate the vascular development, blood pressure, and hypertension-dependent arterial remodeling [18–21]. Recent studies identified Piezo1 as the culprit for VSMC maladaptive mechanosensation in aortic aneurysm, a lethal vascular disease that affects almost exclusively the aged population [22]. Despite its critical role, the downstream effectors of Piezo1 activation that associate with the advance of aging in VSMCs remain unclear. Due to the dynamic nature of the CSK machinery, it plays a critical role in the regulation of mechanosensation of cells to mechanical stimulus [23]. Dysfunctions in VSMC actomyosin architecture, CSK contractile function, and mechanosensation have been demonstrated to cause aging-associated vascular disease such as aortic aneurysm [24]. However, how the changes and interplays between the Piezo1 mechanosensitive channels and cytoskeletal integrity result in pathological mechanosensation in vascular aging still requires further explorations.

Here, we applied a novel ultrasound-tweezers-based micromechanical system to study the changes in VSMCs’ mechanosensation associated with aging. The integrated ultrasound-tweezers system, together the elastic micropillar-based traction force measurement and instantaneous frequency spectrum analysis, would allow us to apply a transient mechanical force to single cell along with temporal analysis of VSMC mechanosensation behaviors. The mechanobiological study revealed that aged VSMCs are associated with a relatively inert solid-like state, resulting in an impairment in their mechanosensation to mechanical perturbations. The decline in mechanosensitive behaviors in aged cells was mediated by hyperactivity of Piezo1-dependent calcium signaling. Determining the molecular drivers of altered mechanosensation behaviors of aged VSMC could provide promising targetable mechanobiological signals that could repress aging-associated cardiovascular dysfunctions and disease progression.

## Results

### Defective VSMC mechanosensation in aging measured with ultrasound tweezers-based micromechanical system

To evaluate the aging-induced defective mechanosensation function of VSMC, we engineered a sophisticated single-cell micromechanical system (**Fig. 1A, Fig. S1**) consisting of an ultrasound tweezers mechanical stress stimulator and an elastic micropillar array force sensor to apply a transient mechanical force to single VSMC and *in-situ* temporal measurements of mechanoresponses to the mechanical perturbation [25–27]. First, polydimethylsiloxane (PDMS)-based micropillar arrays were manufactured using a standard soft lithography technique [28] and used as substrates and cellular force sensors (**Fig. S2;** details see **Methods**). The PDMS micropillar array substrates were microcontact-printed [29] with fibronectin and fluorescence-tagged fibrinogen adhesive proteins on the top of the micropillars (**Fig. S2B**). After cells were cultured on the micropillars, deflections of the micropillars underneath cells were continuously recorded under an inverted fluorescence microscope and used to quantify the forces exerted by the cells (**Fig. 1B, S3**) [30]. Ultrasound tweezers was used to apply a 10-second, 1 Hz, ∼100 pN of transient force to single VSMC through an RGD-integrin bonded lipid-encapsulated microbubble on the cell membrane (**Fig. 1C**, **Fig. S4;** details see **Methods**) under ultrasound excitation, simulating an external mechanical stress [31–33]. In the ultrasound tweezers system (**Fig. S1**), ultrasound pulses generated acoustic radiation force on the microbubble causing its displacement (**Video S1**), and therefore applied controllable mechanical stress on cell.

**Figure 1.**
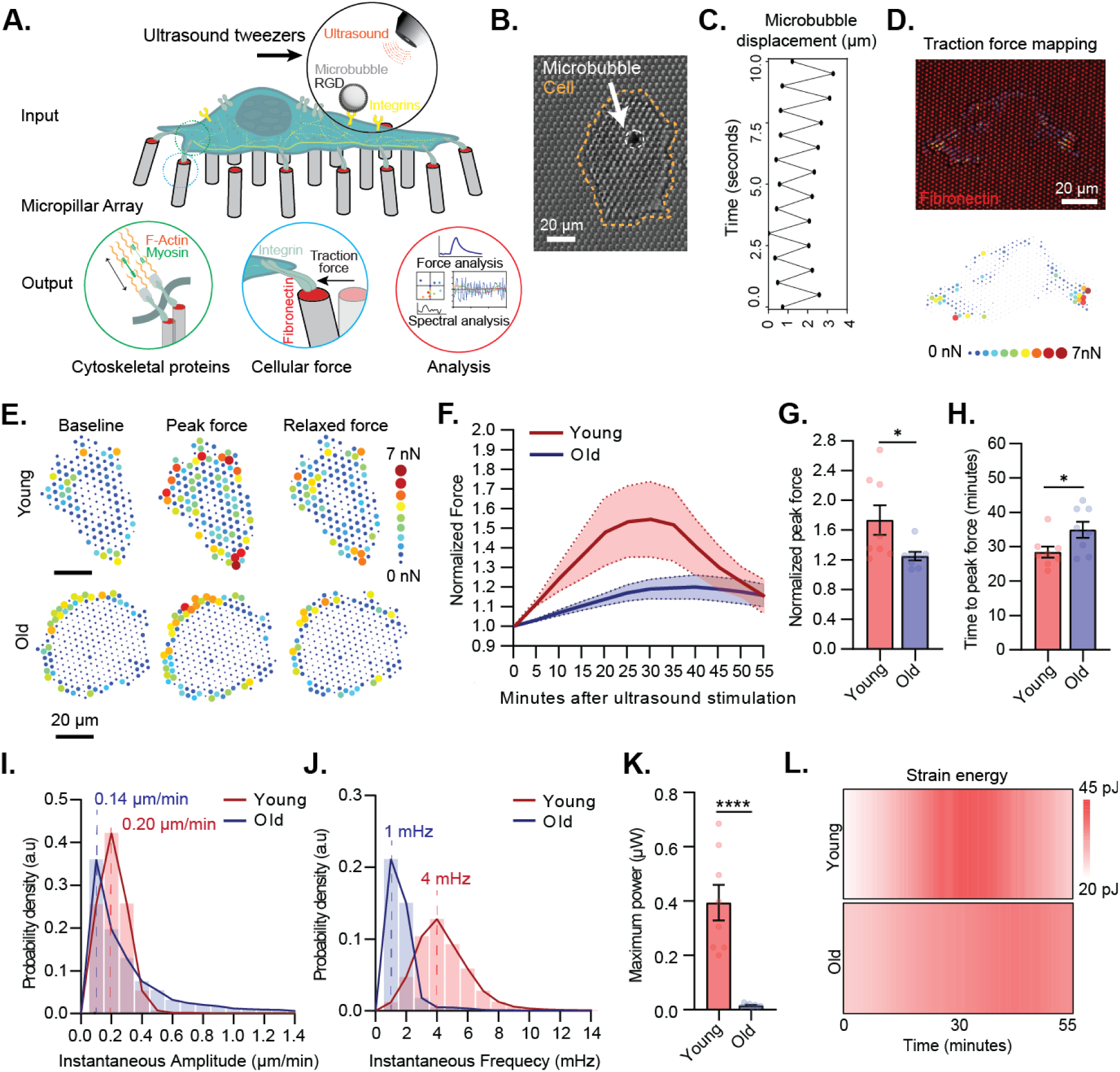
Defective VSMC mechanosensation in aging measured with ultrasound tweezers-based micromechanical system. **A**) A schematic of the integrated single-cell micromechanical measurement system outlining the mechanical input, measurement, and output signals. The ultrasound-tweezers system, together the elastic micropillar-based traction force measurement and force instantaneous frequency spectrum analysis, allow us to apply transient mechanical forces to single cell along with temporal analysis of VSMC mechanosensation behaviors. **B**) A representative bright-field microscopy image showing ultrasound microbubble attachment on VSMC on a PDMS micropillar substrate. VSMC is delineated by dotted orange line, microbubble is shown by white arrow. **C**) Quantified microbubble displacement generated by the ultrasound tweezers under 1 Hz ultrasound stimulation over a period of 10 seconds. **D**) Mapping of traction force of VSMC cultured on a fibronectin-functionalized PDMS micropillar substrate. The top panel shows a representative fluorescent image of micropillar displacements (red channel) and quantified traction force vector map of VSMC (arrows). The bottom panel shows a traction force heatmap. **E)** Representative heatmaps of dynamic traction force evolution of young and old VSMCs in response to a transient mechanical stimulation. Figures show forces pre-stimulation, force during peak contraction and force 10 minutes post peak contraction. **F**) Normalized global traction force in young and old VSMCs evolve after the transient mechanical stimulation. The thick line represents the mean of traction forces over time and the filled area represents the error bar presented as standard error of the mean (SEM). **E**) Normalized peak force and **H**) average time that each cell group took to reach to peak force upon ultrasound tweezers stimulation. **I**) Instantaneous amplitude and **J)** instantaneous frequency distributions of force dynamic responses of young and old VSMCs from 0 to 30 min upon ultrasound tweezers stimulation. The thick lines represent mean amplitude and mean frequency response of young and old cells. **K**) Maximum time-averaged power supplied to single micropillar by young and old VSMCs upon ultrasound tweezer stimulation. **L**) Representative heatmap of average strain energy in young and old cells in response to ultrasound tweezers stimulation. Data presented in F-G are n = 8 cells per group. In G, H, and K, data are presented as mean values ± SEM, P-values were calculated using Student’s t-test. *p <0.05, **p<0.005, ****p<0.0005.

We applied the integrated ultrasound tweezers system to measure mechanosensation responses of primary VSMCs isolated from young (12 weeks) or old (58-78 weeks) mice to a transient mechanical stress on the cell membrane. Following the 10-second exposure to stimulus, for both young and old VSMCs, traction force exhibited a biphasic feature, in which the single-cell level traction force increased within the reinforcement period and gradually recovered in the relaxation period to reach mechano-homeostasis as depicted by the force map of individual cells and quantification of normalized force over the 55-minute observation period (**Fig. 1E-H**). No significant change in force was observed in unstimulated cells (**Fig. S5**). The temporal analysis of force responses revealed that old VSMCs showed compromised ability to generate mechanoallostasic force compared to the young cells (**Fig. 1F**). On average, young VSMCs’ traction force increased 57% compared to the baseline force in the reinforcement period, while the old cells increased only 20% in response to the mechanical stress stimulation (**Fig. 1G**). Meanwhile, old cells not only contracted less, but they also showed a slower and more prolonged contractile phase. The young cells reached their peak contraction within 27.0 ± 2.3 minutes, while the old cells reached the peak contraction in 34.8 ± 6.6 minutes after the stimulation (**Fig 1H**) and failed to reach the basal force level during the 55-minute observation period.

To further delve into the convoluted mechanical signals and analyze the aging-associated shift in VSMC mechanosensation, we performed an instantaneous frequency spectrum analysis based on the single-cell spatiotemporal force measurements, which has been shown to be able to effectively capture sensitive changes in subcellular dynamics across time and space [34, 35]. The force frequency spectrum analysis was based on the Hilbert-Huang transform (HHT), an empirical method that has been successfully applied to explain non-linear and non-stationary signal [36, 37]. We first performed empirical mode decomposition (EMD) to decompose the time series capturing velocity of individual micropillars into a finite and small number of modes called intrinsic mode functions (IMFs) to reduce noise in the original signal (**Fig S6;** details see **Methods**). Application of the HHT to a specified IMF derive the instantaneous amplitude *A*(*t*) and frequency *F*(*t*), so called the instantaneous spectrum that provides frequency magnitude and distribution of the input force dynamics data. Physiologically, the instantaneous amplitude *A(t)* characterizes the magnitude of the force exerted by the cells to move the micropillars, while the instantaneous frequency *F*(*t*) characterizes the rate of which the actomyosin complexes are activated to generate traction force when the cells are subjected to a mechanical stimulation. A higher instantaneous frequency indicates more robust (active) actomyosin CKS dynamics and a higher mechanosensitivity. Thus, the instantaneous spectrum analysis provided a novel approach based on adaptive decomposition of local instantaneous force dynamics of single cells to distinguish spectral difference, and therefore to determine mechanosensation levels among cells.

Upon instantaneous spectrum analysis of young and old VSMCs’ force dynamic responses to the mechanical stimulation, we found that the old cells displayed an abate mechanosensitivity comparing to the young cells, indicated by decreases both in instantaneous amplitude *A*(*t*) (0.15 µm/min) and response frequency *F*(*t*) (3 mHz) (**Fig. 1I, J**). The activation power derived from the instantaneous amplitude and frequency revealed the detrimental effect of aging on VSMC spontaneous force-generating capacity, where the maximum power produced by old cells to move single micropillar were significantly lower than the young cells (**Fig. 1K**). Such changes of VSMC mechanosensation were further specified by generating a temporal heatmap of VSMCs’ strain energy response to reveal the decrease in strain energy consumed by old VSMCs during mechanical maladaptation (**Fig. 1L**) [38]. Altogether, we characterized mechanosensation of VSMCs based on their dynamic force responses and the instantaneous frequency spectrum analyses and revealed a maladaptive mechanosensation in old VSMCs.

### Loss of contractile phenotype and cytoskeletal integrity in VSMCs with aging

The allostatic adaption VSMCs to a mechanical perturbation relies on the interaction of the CSK contractile components to generate cellular force, as well as the engagement of the mechanosensitive signaling elements, i.e. the interplay among mechanosensitive ion channels, integrin-focal adhesion-actin axis, and calcium signal that convert the mechanical input into cellular contractile response [39]. To define the mechanisms underlying aging-associated decline in mechanosensation and VSMC functions, we performed transcriptome analysis of young and old VSMCs using DNA microarray analysis (**Fig 2A**). The transcriptome analysis revealed that 234 and 73 genes had significantly down- or up-regulated expression in old versus young VSMCs, respectively (**Fig. 2B**). To examine how the differentially expressed genes alter functional behaviors of VSMCs in aging, we clustered the differentially expressed genes into different pathways using Gene Ontology (GO) databases. The GO enrichment analysis indicated that several biological processes related to vascular functions were altered with aging (**Fig. 2C,** colored blue; elaborated pathway analysis is included in **Fig. S7**). Enrichment in aging-associated pathways (colored red) confirmed aging phenotype of old VSMCs with enhanced cellular senescence. Moreover, several pathways associated with vascular contractility and mechanosensory behaviors (colored green) were altered in aged VSMCs including vascular smooth muscle contraction, cell adhesion, cell-extracellular matrix (ECM) crosstalk, focal adhesion assembly, regulation of the CSK, detection, and response to mechanical simulation (**Fig. 2C**), supporting the maladaptive mechanosensation in old VSMCs measured by the ultrasound tweezers system.

**Figure 2.**
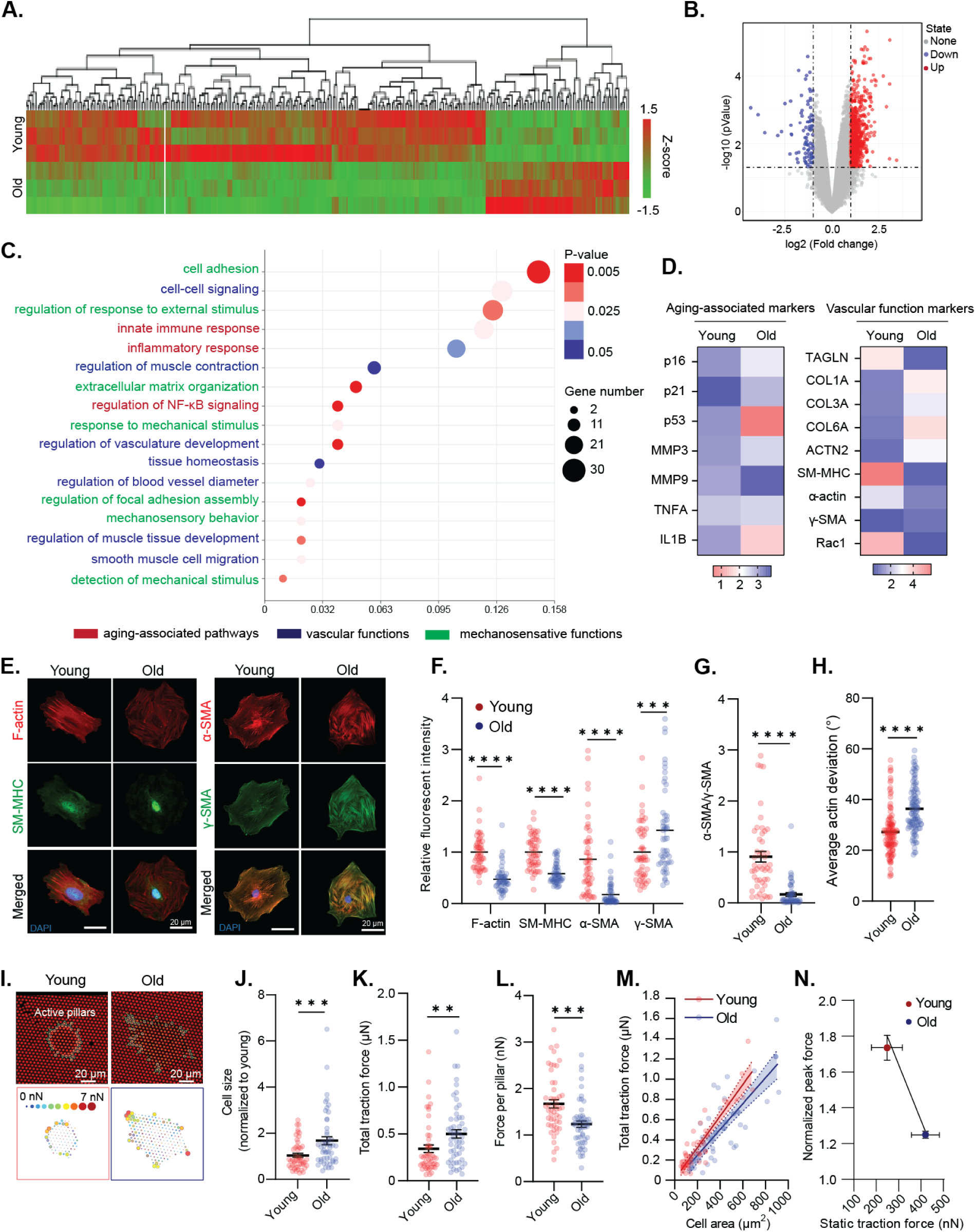
Loss of contractile phenotype and cytoskeletal integrity in VSMCs with aging. **A)** Global gene profiles of young and old VSMCs by DNA microarray analysis (n = 3 for each group). **B**) Volcano plot showing gene expression patterns of all VSMC samples. **C**) Enriched pathways in old VSMC analyzed by GO Enrichment Pathway Analysis. Pathways related to VSMC functional behaviors are highlighted in colors. **D**) qRT-PCR profiled gene expressions related to VSMC functions and senescence in young and aged VSMCs. **E**) Representative immunofluorescence staining images of F-actin, SM-MHC, α-SMA, and γ-SMA in young and old VSMCs as indicated. DAPI is colored blue in merged images. **F**) Quantification of critical cytoskeletal proteins: F-actin, SM-MHC, α-SMA, and ϒ-SMA in young and old VSMCs. Fluorescent intensity of old cell group was normalized to the average intensity of young VSMC group, n = 50 per group. **G**) Ratio of α-SMA/ϒ-SMA showing relative expression of contractile actin isoform over stiffness-maintaining actin isoform in young and old VSMCs (n = 50 per group). **H)** Quantification of the average actin deviation of young and old VSMCs as indicated. n>1000 square regions in both groups. **I**) Representative fluorescent images showing micropillar displacements (top panel). **J**) Quantified cell area, **K**) total traction force, and **L**) traction force per pillar exerted by young and old VSMCs (n = 50 per group). **M**) Regression lines showing correlation between total traction force and cell area of young and old cells, (n = 50 per group). **N**) Correlation between young and old VSMCs’ steady-state total traction force and peak force generated during instantaneous mechanoresponse (n = 8 per group). All data are represented as mean values ± SEM. P-values were calculated using Student’s t-test, *p<0.05, **p<0.005, ***p<0.0005, ****p<0.00005.

To confirm the aging-associated vascular dysfunctions, we profiled gene expression of previously described markers of VSMC functions and senescence, using quantitative reverse transcription PCR (qRT-qPCR) (**Fig. 2D**) [40, 41]. Previous studies have demonstrated increases in cellular senescence, or cell cycle arrest, and senescence-associated inflammatory response with the progress of age [42]. The qRT-PCR analyses showed that genes indicating senescent state such as p16, p21, and p53, as well as inflammation-associated cytokines such as MMP3, IL1B, TNFA were upregulated in old VSMCs. Such changes contributed to VSMC phenotypic switch from a contractile phenotype to a synthetic phenotype and potentiate the risks of cardiovascular disease development [43]. For its significance in VSMC functional behaviors, we performed gene expression analysis on gene encoded for phenotypic plasticity in young and old VSMCs. Healthy VSMCs normally exhibit a differentiated contractile state defined by a high expression of smooth muscle-specific contractile and cytoskeletal genes [44]. However, we found that smooth muscle-specific genes including ACTA2, MYH11, RAC1, and TAGLN were significant downregulated in old VSMCs. Meanwhile, genes indicating synthetic state such as COL3A, MMP1, and MMP9 were strongly upregulated in the old cells. Notably, a strong upregulation of matrix stiffening markers including (COL1A and COL6A) and cytoskeletal stiffening markers ACTG2, ATCN2, and were also observed in these aged cells. The altered gene expression patterns led us to interrogate cytoskeletal remodeling, aberrant mechanosensitive signaling and contractile dysfunction as the culprit of defective mechanoresponse in old VSMCs.

We next assessed the expression and organization of cytoskeletal proteins at the single-cell perspective. We investigated re-organization of the actomyosin CSK with immunostaining and an image recognition-based spatial analysis to characterize partial deviation of the actin stress fibers (**Fig. 2E-H**), in which partial deviation defines the actin alignment uniformity and reflects the bundling and organization of the actin filaments in VSMCs. [45]. Quantitative analysis of CSK architectures indicates that old VSMCs have disrupted actomyosin CSK with lower levels of actin filaments (F-actin) and smooth muscle myosin heavy chain (SM-MHC) (**Fig 2E, F**), along with a higher partial actin deviation (**Fig. 2H & S8**) relative to the young cells, suggesting that the local and total CSK filaments adapted a more diffused and disordered organization with the increase of age. We further explored different smooth muscle-specific actin isoforms and found that aging resulted in a reduction of smooth muscle-alpha actin (α-SMA) and an upregulation of smooth muscle-gamma actin (γ-SMA) (**Fig. 2E, F**). Disruption in α-SMA contractile filaments lead to dysfunction in cytoskeletal tension development in response to stimuli [46], and thereby, is responsible for a decline in VSMC mechanosensation. In addition, γ-SMA in VSMCs resides mainly near the cell cortex and take part in force transmission along the actomyosin-focal adhesion-integrin axis [47]. An increased in γ-SMA has been implied to be a contributor of increased cortical stiffness in previous studies [14, 47, 48]. In aged cells, distribution of γ-SMA stress fiber along the cytoplasm in the absence of α-SMA promoted a stiffening CSK, as indicated by the lower ratio of α-SMA (contractile element) to γ-SMA (stiffening element) in old cells relative to the young cells (**Fig. 2G**).

The measured aging-associated structural alternations in CSK is consistent with the DNA microarray and qRT-PCR analysis results, and suggested aging caused VSMCs transiting from a mechano-active to a more inert solid-like mechano-phenotype, resulting in an impairment in their mechanosensation to mechanical stress. We measured VSMC traction force at their passive, or unstimulated state, where aged VSMCs showed a significant increase in cell area (**Fig. 2J**), exerted a higher total traction force (**Fig. 2K**) but lower average force per pillar compared with that of the young cells, suggesting a shift to a more rigidified phenotype with aging (**Fig. 2J, K**). Further quantitative analysis showed a positive correlation between the total traction force and cell area (**Fig. 2M**). The regression lines were non-parallel, with the slope of the old cells’ regression line were lower compared to the young cells, indicating distinct biomechanical properties between the young and old cells groups. Moreover, while old VSMCs generally have a higher traction force at the passive state, they showed compromised ability to generate mechanoallostasic force compared to the young cells (**Fig. 2N**). This result supported the observations of impaired cytoskeletal integrity and inability of the old cells to generate force during allodynamic process upon mechanical stimulation.

### Piezo1 regulates VSMC mechanosensation via Ca^2+^ signaling and dynamic cytoskeletal remodeling

We next explored the upstream signaling pathway that mediates the decline in VSMC mechanosensation behaviors with aging. Mechanosensation depends on the cell-ECM interaction that promotes conformational changes of the cell membrane and activates the mechanosensitive ion channels in response to mechanical stimuli [49]. In VSMCs, calcium ions have been proven as the “second messengers” molecules that transduce external stimulation into the intracellular cytoskeletal components to trigger actomyosin contractility [50]. Therefore, our next step was to profile gene expression of the common calcium-regulating ion channels [51] in VSMCs to identify the key mediator of VSMC dynamic mechanoresponse. We found that Piezo1, a novel mechanosensitive channel with preference to changes in Ca^2+^ concentration, was remarkably upregulated in old VSMCs (**Fig. 3A**). Immunofluorescence and qRT-PCR results also confirmed that Piezo1 expression was increased by 2 folds in old VSMCs at both mRNA (**Fig. 3B**) and protein levels (**Fig. 3C-D**) relative to the young cells. The Piezo1 ion channel has been known as a key mechanical sensor on the cell membrane regulating Ca^2+^ influx upon its activation driven by external mechanical force and influence cellular force generation since actomyosin contraction is initiated by calcium influx [52–56]. The role of Piezo1 has been previously discussed in cardiovascular disease interventions, including vascular development [43, 44], blood pressure [45], hypertension-dependent arterial remodeling [19], and VSMCs with aneurysm [22], but it remains unexplored in chronological aging. We suspected that the hyperactivation of Piezo1 and Piezo1-mediated calcium signaling are the key mediators of maladaptive mechanosensory behaviors of aged VSMCs.

**Figure 3.**
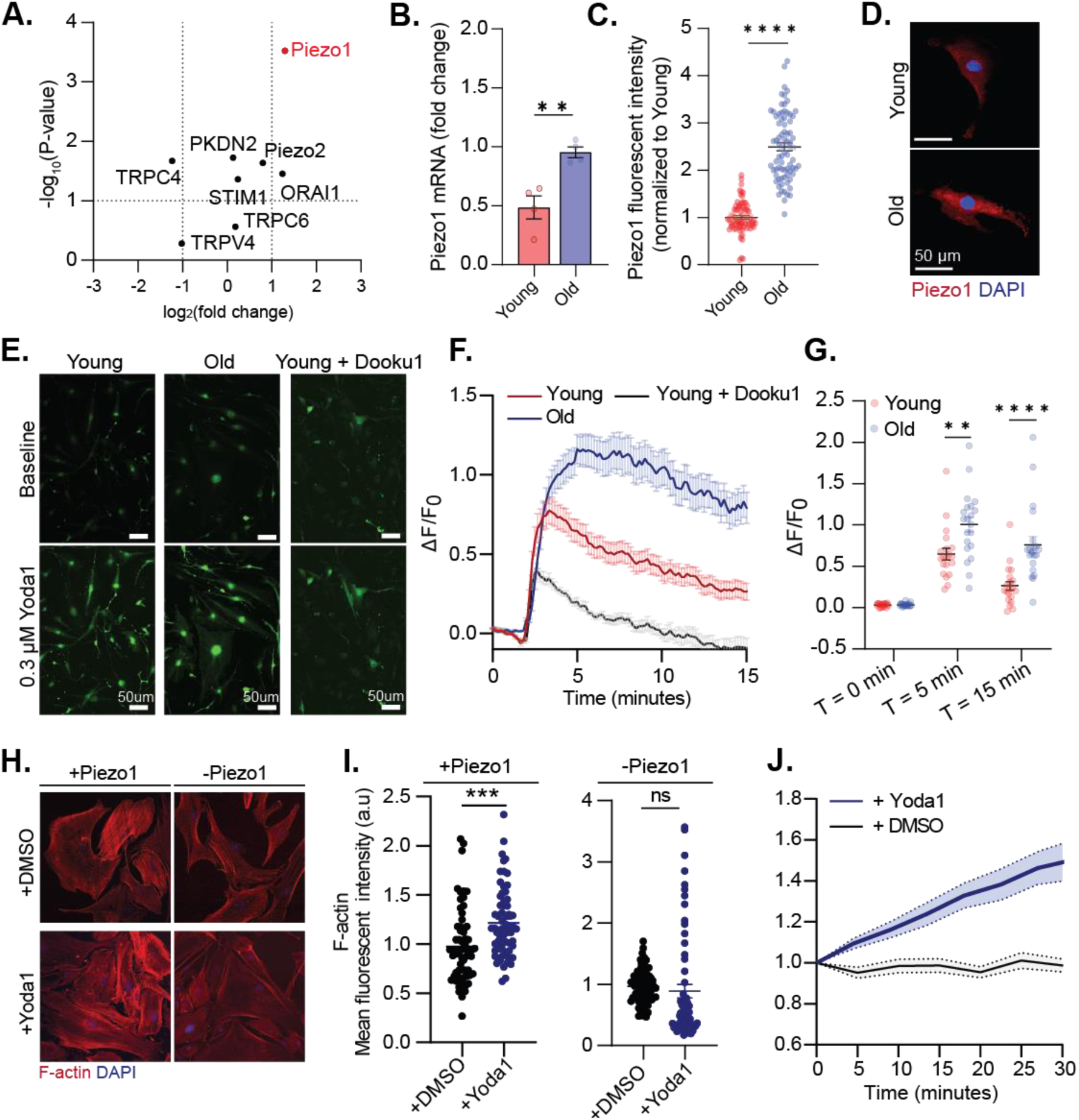
Piezo1 regulates VSMC mechanosensation via Ca^2+^ signaling. (**A**) Dot plot representation showing differential expression (log2) and P value of mechanosensitive channel transcripts in old verses young VSMCs. **B**) qRT-PCR (n = 4 for each group) and **C**) Immunofluorescent analysis of Piezo1 expression in young and old VSMCs (n=50 for each group). **D**) Representative immunofluorescent images of Piezo1 in young and old VSMCs. **E)** Representative images of cytosol Ca^2+^ signal in young, old, and young + Dooku1 cells transfected with Fluo4AM probe. **F**) Time course measurement and **G**) quantification of Ca^2+^ peak response in VSMCs treated as indicated in response to Yoda1 stimulation (n = 20, 20, 5 for young, old, and young+Dooku1 respectively, 3 independent experiments). Intensity of Fluo4AM was normalized to T = 0 min. **H**) Representative immunofluorescent images of F-actin of young VSMCs under different treatment conditions (n = 50 cells per group). +DMSO and +Yoda1 represent VSMCs incubated with DMSO and Yoda1 for 1 hour respectively. +Piezo1 represents control group, and -Piezo1 represents inhibition of Piezo1 activation using Dooku1. **I**) Quantification of F-actin (Phalloidin intensity) of VSMC in indicated groups. **J**) Dynamic force response of young VSMCs in response to Yoda1-mediated Piezo1 activation (n = 5 for each group). + DMSO represents control group. All statistical analysis was performed by Student’s t-test and error bars are presented as SEM. *p < 0.05, **p < 0.005, ****p < 0.0005.

To confirm whether Piezo1 activation determines Ca^2+^ signaling, we transfected the VSMCs with Fluo4 AM, a calcium indicator and quantitatively evaluate changes in cytosolic calcium in response to activation of Piezo1 by Yoda1 – a Piezo1 chemical agonist [57]. Upon addition of 0.3 µM Yoda1, there was an immediate elevation in Ca^2+^ intensity in both young and old VSMCs, followed by a gradual decrease to reach internal homeostasis (**Fig. 3E-F**). Calcium signals were significantly suppressed in the presence of Dooku1, a Yoda1 antagonist that prevents Piezo1 activation, confirming the specificity of calcium influx in VSMCs to Piezo1 channel (**Fig. 3F**). The old cells exhibited a higher peak calcium response and a more sustained calcium signal explained by a more abundant presence of Piezo1 compared to the young cells (**Fig. 3F**). After the 15 minutes observation, calcium signal in the young cells were able to mostly return to the baseline level, while the calcium concentration in the old cells only decreased by 25% (**Fig. 3G**). Altogether, quantification of calcium signaling reinforces that the level of Piezo1 expression determines the persistence of calcium signaling in VSMCs. This conclusion is supported by Pan’s et al findings that Piezo1 triggers a biphasic response of intracellular calcium, where overexpression of Piezo1 causes a prolonged calcium signaling and disruption in the actomyosin-focal adhesion-ECM pathway [58]. This calcium patterns reflected the biphasic mode of VSMC allostasis force dynamic measured by the ultrasound tweezers system (**Fig. 1G**), suggesting that Piezo1-dependent calcium signaling regulates the persistence of the mechanoresponse in VSMCs, while the activation level of contraction is dependent on the concentration of contractile filaments presented in the CSK.

Ca^2+^ influx is an upstream signal that initiates tension development in the CSK via cooperation of the actomyosin complex [59]. The natural next step we took was to interrogate Piezo1 as the long-range mediator for actin remodeling and force generation in young (healthy) VSMCs. We performed quantitative assessment of F-actin intensity after 1-hour of Yoda1 treatment using phalloidin staining. Yoda1-induced Piezo1 activation resulted in an elevation of F-actin expression compared to their basal state (**Fig. 3H, I**). Conversely, no significant change was detected in cells treated with Dooku1 in prior to treatment of Yoda1. These findings revealed that opening of Piezo1 mechanosensitive channel is essential for actin polymerization in response to extracellular stimuli. Next, we measured Yoda1-stimulated changes in CSK tension of VSMC using the micropillar array and confirmed that Piezo1 activation led to an increase in force response of VSMCs (**Fig 3J**). Altogether, our results identified Piezo1 as a critical regulator of calcium signaling and cytoskeletal force development in VSMCs and suggested that overexpression of Piezo1 is the upstream culprit of the maladaptive mechanoresponse in old VSMCs.

### Piezo1 expression can be fine-tuned by *in vitro* aging models

Based on the experimental results above, we asked whether extenuating Piezo1 overexpression would mitigate the loss in mechanosensitivity of the old cells. We therefore established two *in vitro* models that aimed to fine-tune Piezo1 expression to regulate mechanical aging of VSMCs. In the first model coined “reversed aging”, we performed siRNA transfection with ON-TARGETplus siPiezo1 at 25 nM for 48 hours to reduce Piezo1 at mRNA level in old VSMCs. The second model called “induced aging” involves acceleration of aging in young VSMCs by pharmacological treatment of 10^-7^ M Angiotensin II (AngII) for 48 hours. AngII has been found to provoke vasoconstriction [60], inflammation [61], and vascular remodeling [62], thus lead to cardiovascular disease development, such as atherosclerosis and hypertension [63]. Kunieda et al discovered that AngII induced premature cellular senescence of human VSMCs via a p21-dependent pathway [64]. Our gene expression analysis revealed a strong correlation between aging and cellular senescence, where old cells showed upregulation of cell cycle proteins, notably p21, (**Fig. 2D**). Thus, cell senescence-driven aging by AngII would be an efficient *in vitro* model to mimic the effects of chronological aging on VSMC Piezo1-dependent mechanosensation.

To elaborate on VSMC heterogeneity and functions among the regulated aging pathways, we performed transcriptome profiling using single-cell RNA sequencing. Single-cell suspensions were prepared and sequenced for AngII-treated (induced aging) young cells and the siPiezo1-treated (reversed aging) old cells, as well as the control young and old cells (**Fig. 4A**). The clustering analysis captured three distinctive regions among the sequenced cells. Based on the experimental conditions of the input, we were able to distinguish the young, old, and AngII-treated young cell groups, while the siPiezo1-treated old cells exhibited significant overlap with the cluster of old cells (**Fig. 4B** & **Fig. S9**). The young cells were characterized by abundant expression of VSMC markers, such as α-actin2 (Acta2) and Myosin light chain 6 (Myl6), while the old and AngII-treated young cell groups were classified by a significant upregulation of aging-associated genes. Particularly, we found that genes related to cellular senescence (e.g., Ccdn1, Ccdn2, Ets1, Ptgs2, Ilrl1) and senescence-associated secretory phenotype (Mmp3) were highly concentrated in the clusters of old and AngII-treated young cells (**Fig. 4C, D**). The enriched expression of Mmp3 in VSMCs were previously shown to be regulated by Piezo1 activation in VSMC of aortic aneurysm, suggesting that the old and AngII-treated young cells are associated with dysfunction in mechanosensory behaviors and high risk of cardiovascular disease development [22]. Notably, we found that the AngII-treated young cell cluster showed the highest amount of CD44 (macrophage-like VSMC marker) among the four groups, while SPP1 (osteogenic VSMC marker) was only found in the old and siPiezo1-treated old cell groups (**Fig. S10**). Based on previous reports of the multi-directional phenotypic switch in VSMCs in response to different extrinsic signal, we hypothesized that VSMC transformed into osteogenic state during chronological aging and macrophage-like state in AngII-induced aging [65, 66]. Interestingly, despite the considerable difference between the AngII-treated young cell and the old cell clusters, we found a resemblance in the differential expression pattern of the cytoskeletal markers between the old and AngII-treated young cell (induced aging) groups. By profiling the contractile genes such as α-actin2 (Acta2), γ-actin (Actg1), Tagln (Transgelin), and α-actinin1 (Actn1), we confirmed that AngII treatment is efficient in mimicking the effect of vascular aging on VSMC cytoskeletal integrity. The single-cell gene profile between young and young AngII group showed a downregulation of α-actin2, in contrast to an upregulation of γ-actin, transgelin, and α-actinin. The results resembled our qRT-PCR and immunostaining results measuring changes in contractile gene expression with normal aging, as shown in **Fig. 2D-F**. Thus, it is suggested that pharmacological treatment of young cells with AngII disrupted the CSK-mediated contractile properties in young cells and promoted their transition into a more mechanically passive state, while siPiezo1 treatment does not alter the CSK integrity and the baseline mechanical behaviors of old cells.

**Figure 4.**
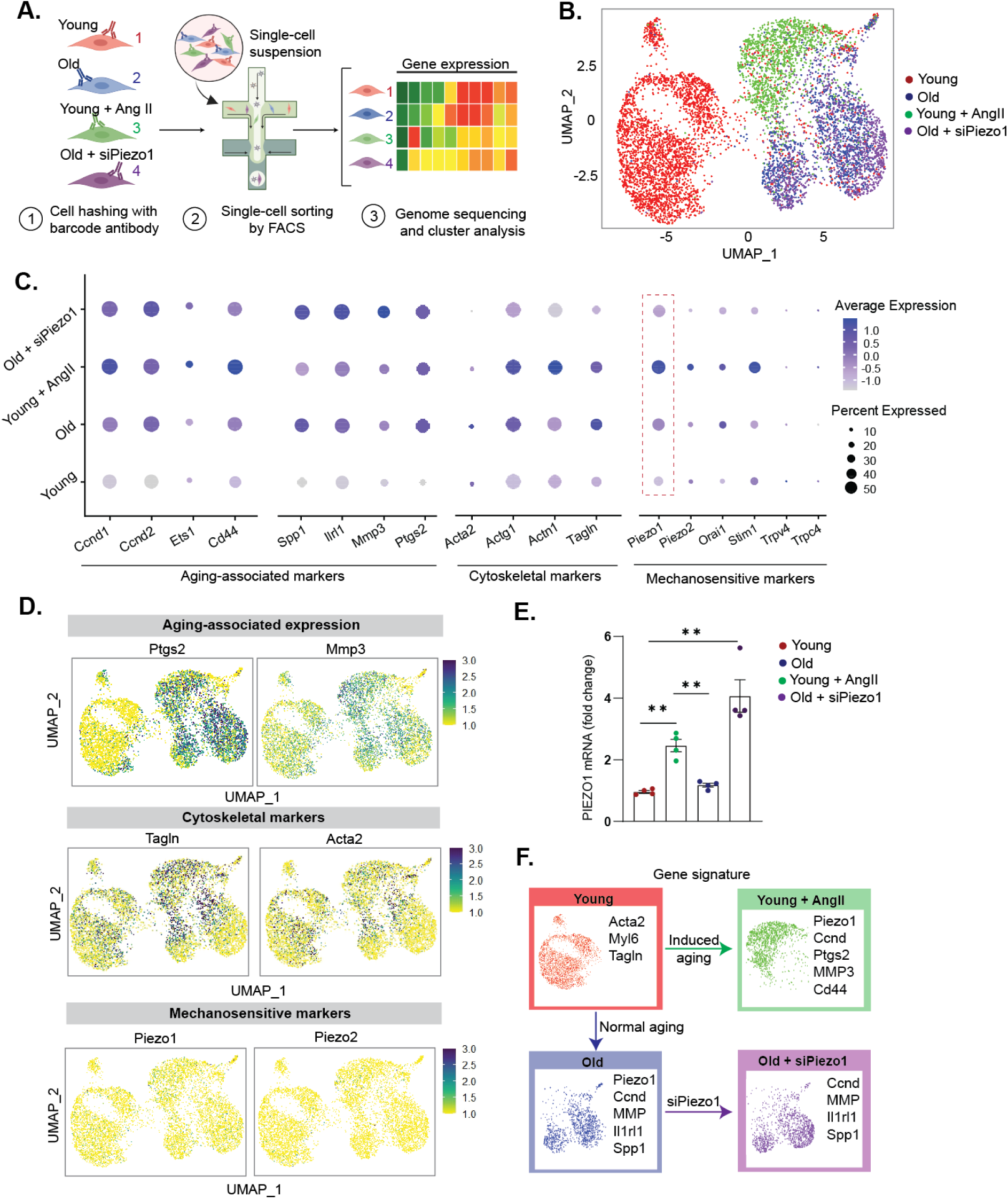
Single-cell sequencing analysis of *in vitro* aging models. **A**) Schematic showing study overview. **B**) UMAP (uniform manifold approximation and projection) visualization of color-coded clustering of the VSMC in the four experimental groups (young, young + AngII, old, and old + AngII). **C**) Dot plot showing average and percent expression of genes encoded for aging-associated markers, cytoskeletal markers, and mechanosensitive markers in the four cell clusters. **D**) Representative UMAP showing expression levels of classified genes. Two representative genes from each indicated group were presented. **E**) qRT-PCR analysis of Piezo1 expression in VSMCs of young, young + AngII, old, and old + AngII respectively (n=4 for each group). All data are represented as mean values ± SEM. **F)** Summary showing upregulated genes in each cell cluster. **p<0.005. Figure 4A is created with BioRender.com.

Next, we examined the mechanosensing receptors presented in the four cell clusters. Among the mechanosensitive channels, Piezo1 was highly expressed and was positively associated with aging pathway, indicated in both old and AngII-treated young cell (induced aging) groups (**Fig 4C-D**). Importantly, siPiezo1 treatment significantly reduced the Piezo1 expression level in the old cells, confirming successful siRNA transfection. We further validated the results using qRT-PCR and found that Piezo1 mRNA level was alleviated by 1.5 folds in the siPiezo1-treated old cells compared to the old cells, while there is no significant difference between the siPiezo1-treated old cells and young cells. This result suggested that overexpression of Piezo1 is the hallmark of aging and aging-associated cardiovascular disease in VSMCs. From the single-cell analysis of upregulated genes in each experimental cluster, we confirmed that while AngII treatment induced vascular aging in a different pathway compared to chronological aging, it provided a considerably efficient method to regulate aging *in vitro* by mimicking the changes in contractile functions and mechanosensitive Piezo1 channels. Meanwhile, siPiezo1 transfection is good for specific targeting of Piezo1 expression to fine-tune the mechanosensitive signaling pathway without shifting the phenotypic signature of the old VSMCs (**Fig. 4F**).

### Antagonizing Piezo1 mitigates aging-mediated aberrant mechanosensation in old VSMCs

To further evaluate Piezo1 mechanosensitive signaling in the established i*n vitro* models (**Fig. 5A**), we assessed Piezo1-mediated calcium signaling and mechanoresponse of VSMCs in each group upon Piezo1 activation. We first performed immunostaining to detect the protein expression level of Piezo1 and found that Piezo1 expression was alleviated 1.8 folds at protein levels in the “reversed aging” cells compared to the old cells (treated with non-targeting siRNA). In the “induced aging” model, Ang II-treated young cells showed an increase in Piezo1 by 2 folds compared to the control young cells (**Fig. 5B**). We next examined whether manipulating Piezo1 expression in the two designed cell groups can mimic the calcium signaling patterns that we observed in VSMCs during chronological aging by applying Yoda1 chemical stimulation. In response to Yoda1 addition, both the siPiezo1-treated (reversed aging) old cells and the AngII-treated (induced aging) young cells exhibited a rise in cytosolic Ca^2+^ compared to their basal state, with a higher calcium peak observed in the induced aging cells. 10 minutes after treatment of Yoda1, the reversed aging cells were able to achieve a homeostasis condition, while the induced aging cells fails to return to its baseline calcium level (**Fig. 5C**). The results not only validated the efficiency of pharmacological treatments to simulate the mechanical aging process *in vitro*, but also confirmed the regulatory roles of Piezo1 on calcium signaling in VSMCs. Furthermore, silencing Piezo1 prompted the promising approach to enervate the hyperactivation of Piezo1 in old cells while ameliorated their ability to adapt rapidly to an external stimulation in calcium-related niche. Next, we investigated actin CSK and static force of VSMCs in the reversed aging and induced aging models. Inhibition of Piezo1 in old cells had no significant effects on their F-actin intensity, implicating that Piezo1 plays a less important role on the CSK at the non-dynamic level (**Fig. 5D-E**). Meanwhile, AngII treatment resulted in a decrease by 2.4 folds in F-actin concentration in young cells, therefore perturbing their ability to generate dynamic force in response to a mechanical stimulation. Overall, the induced aging group using AngII treatment demonstrated a total shift in the mechanophenotype of young VSMCs, including changes in actin CSK, cell area, and static traction force while the reversed aging group using siPiezo1 treatment conserve the biophysical properties of the old cells at the baseline perspective (**Fig. 5F**).

**Figure 5.**
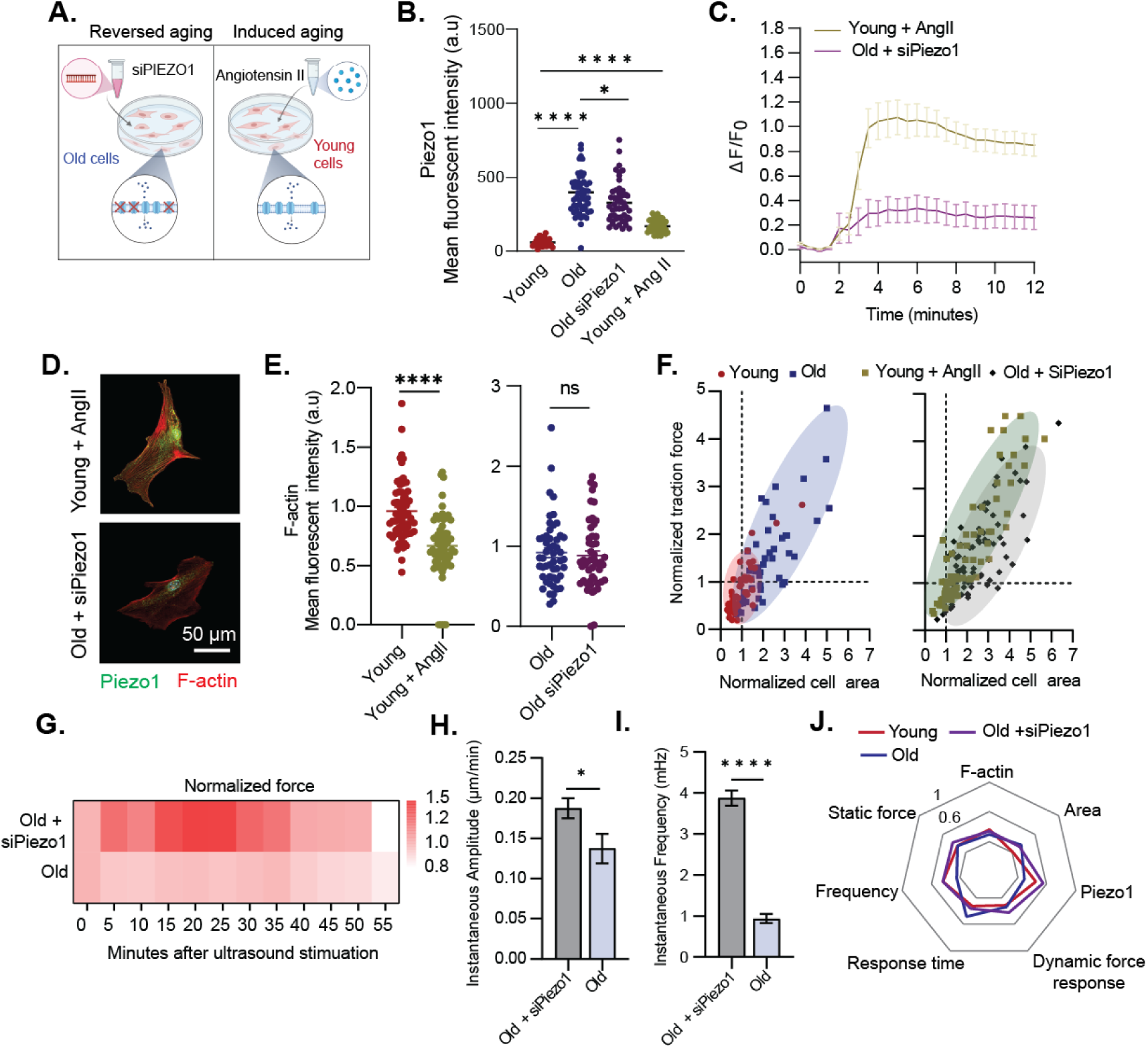
Fine-tuning Piezo1 recovered mechanosensation in old VSMCs. Fine-tuning Piezo1 recovered mechanosensation in old VSMCs. **A**) Schematic showing two *in vitro* models using pharmacological treatments to mimic biological aging and regulate Piezo1 expression. **B**) IF analysis of Piezo1 expression in VSMCs of young, young + AngII, old, and old + AngII respectively (n=50 for each group). **C**) Time course of Ca^2+^ signal in VSMCs treated as indicated in response to Yoda1 stimulation (n = 20 for each group, 3 independent experiments). Intensity of Fluo4AM was normalized to T= 0min. **D)** Representative IF images of Piezo1 and F-actin fluorescent expression in young + AngII and old + siPiezo1 VSMCs**. E**) Quantification of changes in F-actin expression (Phalloidin) in young versus young + AngII VSMCs and old versus old + siPiezo1 VSMCs (n=50 for each group). Fluorescent intensity of young versus young + AngII group was normalized to young, and fluorescent intensity in old versus old + siPiezo1 group was normalized to old cells. **F**) Clustering diagrams showing the correlation between cell area and traction force at static state in all VSMC groups treated as indicated (n=50 for each group). **G**) Temporal heatmap of force response of old cells treated with siPiezo1 and old cells upon ultrasound stimulation (n=8 cells), results are normalized to average force response of old cells. **H**) Instantaneous amplitude and **I**) Instantaneous frequency of old versus old-siPiezo1 cells (n =8 cells). **J**) Representative radar chart showing the average values of mechanical phenotypes of VSMCs in different group. Data are normalized between 0 and 1 using min-max normalization. All data are represented as mean values ± SEM. * denotes p<0.05, **p<0.005, ***p<0.0005, ****p<0.00005. Figure 5A is created with BioRender.com.

Building upon these findings, we next measured whether reducing Piezo1 overexpression aid in recovery of VSMC mechanosensation by applying the ultrasound tweezers system measurement to probe force response of old cells treated with siPiezo1. Upon a 10-second transient stimulation, we observed that the old cells treated with siPiezo1 exhibited a higher mechanosensitivity, as demonstrated by a stronger increase in cellular force compared to the control old cells (**Fig. 5G**). The treated cells also recovered their biphasic mechano-allostasis behavior with an excitation period from 15 min to 37.5 min, followed by a gradual decrease in force for the remaining 17.5 mins. We then applied HHT-based spectral analysis to examine the instantaneous characteristics of the two cells group through their temporal force response. We found that the old cells treated with siPiezo1 showed a significantly higher instantaneous amplitude compared to the control old cells during the 55-minute period (**Fig. 5H**), as well as a higher instantaneous frequency distribution compared to the old cells with no treatment (**Fig. 5I**). This shift in instantaneous frequency spectra indicated that inhibiting overexpression of Piezo1 in old cells improved their force-generating capacity as well as the dysfunctions in mechanosensitive signaling, thereby restored their ability to adapt rapidly to external mechanical stimulation. Overall, antagonizing Piezo1 expression considerably restored the old cells’ mechanosensing properties to resemble to that of the young cells (**Fig. 5J**). Ours results highlighted the tremendous efficiency of fine-tuning Piezo1 in regulating the mechanosensitive signaling pathway underlying VSMC mechanoresponse and suggested that suppressing Piezo1 expression would provide a therapeutic treatment to partially ameliorate the biophysical loss caused by aging in VSMCs.

## Discussion

The dynamic interplay between the mechanosensitive receptors and the intracellular structures allows VSMCs to alters their physiological and biophysical behaviors to rapidly adapt to the mechanical perturbations in the microenvironment and maintain an internal homeostasis. Aging leads to detrimental disruptions in VSMCs’ mechanosensitive behaviors from the cellular and subcellular levels. Substantial studies regard disruptions in the CSK and cell-ECM interaction with aging, eventually leading to dysfunctions in the force-generating capacity during mechanical adaption, have been reported [67, 68]. Understanding of aging-altered mechanosensation arising from single-cell perspective is necessary to elucidate the dynamic molecular processes underlying cardiovascular disease development in the elderly population. However, due to the lack of proper spatiotemporal cellular force measurement and analysis tools, most studies have been limited to studying cells at the colonial and static levels. Herein, we developed a unique ultrasound tweezers system that enables manipulation over the mechanical force applied to single cells, *in situ* assessment of cellular force dynamics (e.g., response time and contractile magnitude), and subsequent advanced signal processing (e.g., instantaneous force frequency analysis) for in-depth analysis of VSMC intrinsic mechanosensation behaviors.

Using the ultrasound tweezers system, we explored the compromised mechanosensation regard a decline in contractile force and a delayed force response time in old VSMCs after exposure to a transient mechanical stimulation. The results point toward a common biomechanical pattern demonstrated in our previous findings where a blunt mechanosensation was found in VSMCs with progressive aortic aneurysm, therefore extending the understanding on VSMCs’ force dynamics from a pathological condition (aneurysm) to a non-pathological context (chronological aging) [22]. The maladaptive mechanosensation were furthered characterized by a novel signal processing framework based on an advanced instantaneous force frequency spectrum analysis method that provided instantaneous amplitude and instantaneous frequency to the time-varying force measured by the ultrasound tweezers system. The instantaneous force frequency spectrum helped us delve deeper into understanding the biophysical nature of the input force signal and intrinsic changes in the cytoskeletal structures that drive VSMC mechanoresponse. We showed that aging in VSMCs caused a reduction in their spontaneous force-generating power, as well as in the dynamic interaction among the actomyosin complexes, featured by a negative shift in both instantaneous amplitude and instantaneous frequency.

Such decline in mechanosensation were further explained by the changes in transcriptome signatures of VSMCs with aging. Based on a DNA microarray array analysis, we found that old VSMCs were strongly correlated with cellular senescence and inflammatory responses, as well as significant alterations in the pathways associated with vascular functions and mechanosensation. qRT-PCR results targeting specific expression of individual gene expression also showed that aging causing a phenotypic switch of VSMC from a contractile phenotype to a synthetic phenotype, leading to defective contractile functions and stiffening of the CSK. At both genetic and protein levels, we found a significant alleviation of VSMC contractile markers in old VSMCs (e.g., F-actin, smooth muscle-specific α-actin, and myosin) and an upregulation of cytoskeletal stiffening markers (γ-actin). Notably, other studies by Massett *et al* and Seawright *et al* also emphasized the increased of γ-actin throughout in the loss of α-actin as the subcellular culprit behind the decline in contractile functions, mechanosensing, and cell-matrix adhesion in VSMCs [14, 46]. Adding on to such findings, our study used the ratio of α-actin over γ-actin as the complementary “stiffening index” to characterize stiffening of the VSMC CSK with aging. Furthermore, the disruption in cytoskeletal integrity and organization altered traction force of old VSMCs at both static and dynamic level. At the basal state, we found that despite the increase in the total traction force in old VSMCs, the average force per pillar exerted by the old cells were significantly lower compared to the young cells, thus indicated an impairment in the contractile force ability that drives aberrant VSMC mechanosensation.

This study also uncovered the role of Piezo1 mechanosensitive signaling in aging-dependent mechanobiological dysfunctions (**Fig. S11**). We first discovered a substantial elevation in Piezo1 expression in old VSMCs at both transcriptome and proteome levels, which was a pathological hallmark indicated in previous studies on vascular cells with pulmonary arterial hypertension [18] and cardiomyopathy [69]. We revealed that the overexpression of Piezo1 in old VSMCs mediated abnormal calcium signaling and altered the downstream pathway associated with calcium influx. Activation of Piezo1 in intracellular calcium initiates the sliding of actomyosin myosin complexes and resulting in an increase in cellular force [70]. In normal young cells, calcium influx is transient upon Piezo1 activation, while in the old cells, the increased calcium concentration is sustained for a prolonged period. We predicted that the rise in intracellular calcium level in the old cells exceed the physiological requirement within VSMCs, thereby induce vascular dysfunctions. With increasing age, the regulatory ability of cytosolic Ca^2+^ attenuates, disrupting the signal transduction within the ECM-integrin-cytoskeletal axis regulating VSMC contractility and making VSMC to be stiffer and less sensitive to mechanical stimulation.

Building upon our findings, we provided a potential “mechanomedicine” strategy aiming to reduce Piezo1 overactivation in aged VSMC to restore a healthy mechanosensation, and thus alleviate the mechanical disruptions in vascular aging. To examine whether Piezo1 expression can be manipulated *in vitro*, we designed two models utilizing pharmacological treatments to induce “premature aging” using AngII and “reverse aging” by antagonizing Piezo1 expression with siRNA transfection. To examine the effects of those treatment on the mechanophenotype of VSMCs, we performed whole transcriptomic profiling using single-cell sequencing analysis to detect expression of targeted aging-associated, cytoskeletal, and mechanosensitive markers in the four experimental groups: young, old, induced aging, and reverse aging. The gene signature in each gene cluster indicated that AngII induced aging in young cells can partially mimic *in vivo* chronological aging, supporting by the similar differential gene patterns with that of the old cells measured by quantitative PCR and immunofluorescence. Meanwhile, siRNA transfection was efficient to antagonize Piezo1 overexpression without altering the transcriptomic signature. We tested Piezo1-mediated signaling in both models and confirmed that AngII-treated young cells exhibited an impairment in calcium signaling, thus induced dysfunctional mechanosensitive signaling that was found in the normal old cells. Conversely, reducing Piezo1 overexpression in the old cells by siRNA transfection ameliorate their mechanosensation, confirming by a recover in instantaneous frequency response upon ultrasound tweezers stimulation.

Taken together, combining the ultrasound tweezers micromechanical system, transcriptome profiling (DNA microarray and single-cell RNA sequencing), and advanced mechanobiological analytic methods, we established a complete framework for the analysis of VSMC mechanosensation behaviors and characterizations of the mechanosensation dysfunction associated with aging. We established the paradigm in which abnormal expression of mechanosensitive Piezo1 ion channel plays a long-range regulatory role in aging-associated impairment in VSMC mechanosensation. This opens a world of possibilities for broader applications, that are not only limited to vascular cells, but to any adherent cell, and across a multitude of problem areas such as determining etiology of diseases like cardiovascular diseases, diabetes, and Alzheimer’s; epithelial-to-mesenchymal transition that occurs during cancer tumorigenesis, angiogenesis and vasculogenesis and many more.

## Methods

### Cell culture

Young and old mouse primary aortic VSMCs (CSC-C4696L and CSC-C4301X, purchased from Creative Bioarrays) were isolated from aortas of C57BL/6 mice of 12 weeks and 58-78 weeks, and were cultured in SuperCult® Mouse Aortic Smooth Muscle Cell Medium Kit (CM-4696L and CM-C4301X, Creative Bioarrays). Cell culture media was replaced every 2 days. For all experiments, primary VSMCs were cultured for no more than 4 passages.

### Micropillar array fabrication

The PDMS micropillar array fabrication was carried out using a standard soft lithography two-step molding process [28]. A silicon master mold (micropillar height 7.1 µm and diameter 1.8 µm) was fabricated using photolithography and deep reactive ion etching. Silanization of silicon molds was carried out with tridecafluoro-1,1,2,2, -tetrahydrooctyl)-1-trichlorosilane (Sigma-Aldrich) overnight in vacuum. Negative PDMS molds were made from the silicon master mold by adding Sylgard 184 silicone elastomer base and curing agent (Dow Corning) mixture by 10: 1 mass ratio to the silicon master mold, and baking at 110°C over 30 mins. After curing, the negative PDMS molds were peeled off from the silicon mold and silanizied in vacuum overnight. PDMS micropillar arrays were generated by casting a layer of Sylgard 184 silicone elastomer base and curing agent mixture by 10: 1 mass ratio on the surface of the silanized negative molds, and the negative molds were then covered with oxygen plasma (350 W, PlasmaEtch) treated 22mm x 22 mm glass coverslips (Electron Microscopy). After curing in an oven heated at 110 °C for 48 hours, the glass coverslips containing PDMS micropillar arrays were peeled off from the negative molds. The PDMS micropillar arrays were immersed in 100% ethanol and sonicated to restore collapsed micropillars, then dried with a critical point dryer (Samdri®-PVT-3D). The glass coverslips containing PDMS micropillar arrays were mounted on a 60 mm petri dish with a 15 mm hole in the center. Finally, the PDMS micropillar arrays were functionalized using microcontact printing with fibronectin (50 μg/ml; Sigma-Aldrich) and Alexa-Fluor 647-conjugated fibrinogen (25 μg/ml; Life Technologies). Schematic illustration of micropillar fabrication and functionalization is presented in **Fig. S2**. VSMCs were then seeded on the functionalized PDMS micropillar array in the petri dish and cultured overnight prior to experimentation.

### Microbubble attachment and mechanical stimulation of VSMCs using ultrasound tweezers

Biotinylated VesselVue microbubbles (Sonovol) of diameter between 4-5 µm were used in this study. First, the microbubble solution was mixed with streptavidin (10 mg/ml, ThermoFisher) at a volume ratio of 20:1 for 30 min and kept at room temperature to form streptavidin-conjugated microbubbles via streptavidin-biotin binding. The streptavidin-conjugated microbubbles were washed twice with PBS to remove the unbound streptavidin. Next, biotinylated Arg-Gly-Asp (RGD) peptides (2 mg/ml, Vividtide) were added to the streptavidin-coated microbubble solution in the ratio of 2:21 for 20 min at room temperature, then 1.2 μL of the microbubble solution was collected from the top layer and mixed with 48.8 μL of cell culture medium. Cell culture medium was aspirated from the petri dish containing VSMCs on micropillars. The prepared microbubble solution was added on top of the micropillar arrays with adherent cells. The petri dish was then flipped for 10 min to allow microbubbles to bind to the adherent cells via floatation. Lastly, the petri dish was flipped back and gently washed to remove any unbound microbubbles.

VSMCs with only 1-2 microbubbles attached were selected for experimentation with ultrasound tweezers. A 10-MHz ultrasound transducer (V312-SM, Olympus) was used to apply acoustic ultrasound pulses to move the microbubbles so as to stretch the cell membrane at a frequency of 1 Hz for a period of 10 s. Before the experiments, the ultrasound transducer was aligned using a pulser receiver (Olympus) to fix at a 45° angle and a 11.25 mm distance (Rayleigh distance) from the target cell under an inverted microscope (Zeiss Axio Observer Z1). A function generator (Agilent Technologies 33250 A) along with a 75-W power amplifier (Amplifier Research 75A250) was used to drive the ultrasound transducer. After the ultrasound stimulation, deflections of the micropillars were recorded continuously for one hour with a 30-s interval with the microscope, and traction force exerted by VSMCs during dynamic mechanical response was assessed and quantified as described above.

### Traction force measurement and spectrum analysis of VSMC force dynamics

After the 1-Hz, 10-s ultrasound stimulation, deflections of individual micropillars underneath cells were recorded continuously for one hour with a 30-s interval with an inverted microscope (Zeiss Axio Observer Z1), and traction force exerted by VSMCs at each time point during the dynamic response was quantified based on the displacements of the fluorescence-tagged micropillars using Cellogram and customized MATLAB programs (Mathworks) (details see **Fig. S3**) as described previously [22, 30]. The instantaneous force frequency spectrum analysis was obtained by tracking the time series of displacement velocity of individual micropillars following the ultrasound stimulation. Specifically, EMD was performed separates the velocity time series into a finite and small number of IMFs (a portion of the complete signal) to reduce noise in the original signal. HHT was then applied to each IMF to derive instantaneous frequency F(t) and instantaneous amplitude A(t) at each time point (t) [22, 34], and generate the instantaneous spectrum that provides frequency magnitude and distribution of the input force dynamics data. In this study, IMF2 was used to calculate the instantaneous characteristics of VSMC force response due to its optimal noise reduction and resemblance with the original signal.

### Immunofluorescence staining and microscopy

VSMCs were fixed in a 4% Paraformaldehyde (Alfa Aesar) solution for 20 minutes and permeabilized with 0.2% Triton X-100 (Roche Applied Science) in phosphate buffered saline (PBS, Gibco) for 10 minutes. Then, VSMCs were incubated in 3% bovine serum albumin (BSA, Sigma Aldrich) blocking buffer for 1 hour at room temperature to prevent nonspecific bindings. Staining was performed by incubating the cells overnight at 4°C with primary antibodies diluted in 1% BSA targeting α-smooth muscle actin (1:200; 48938S, Cell Signaling Technology), gamma actin (1:500; PA1-16890, Invitrogen), smooth muscle myosin heavy chain (1:200; PA5-116524, Invitrogen), Piezo1 (1:25; NBP178446, Novus Biologicals), alpha-actinin 2 (1:100; MA531539, Invitrogen). Primary antibodies were then removed, and samples were washed 4 times with PBS with Tween 20 (PBST, ThermoScientific), followed by application of diluted secondary antibodies in 1% BSA for 1 hour at room temperature. Alexa Fluor 647-conjugated anti-rabbit IgG (1:500; 4414S Cell Signaling Technology) and Alexa Fluor 488-conjugated anti-mouse IgG (1:500; SAB4600066, Invitrogen) were used as secondary antibodies for fluorescent signal detection. Alexa Fluor 555-conjugated Phalloidin (1:300; A34055, Invitrogen) were used for visualization of F-actin filaments. The cells were washed 4 times with PBST and counterstained with 4′,6-diamidino-2-phenylindole (DAPI; Invitrogen) at 1:1000 dilution. After immunofluorescence staining, images were captured using a Zeiss LSM 710 confocal microscope (Carl Zeiss) and acquired via the Zeiss Efficient Navigation (ZEN) software (Carl Zeiss).

### Quantitative real-time PCR (qRT-PCR)

RNA isolation was performed using the Directzol RNA MiniPrep Kit (R2052, Zymo Research). Following reverse transcription using the cDNA Synthesis Kit (1708890, Bio-Rad), quantitative real-time PCR was performed with Fast SYBR Green Master Mix (4385610, Life Technologies) using a CFX96 Touch Real-Time PCR Detection System (BioRad). The primer sequence set used were:

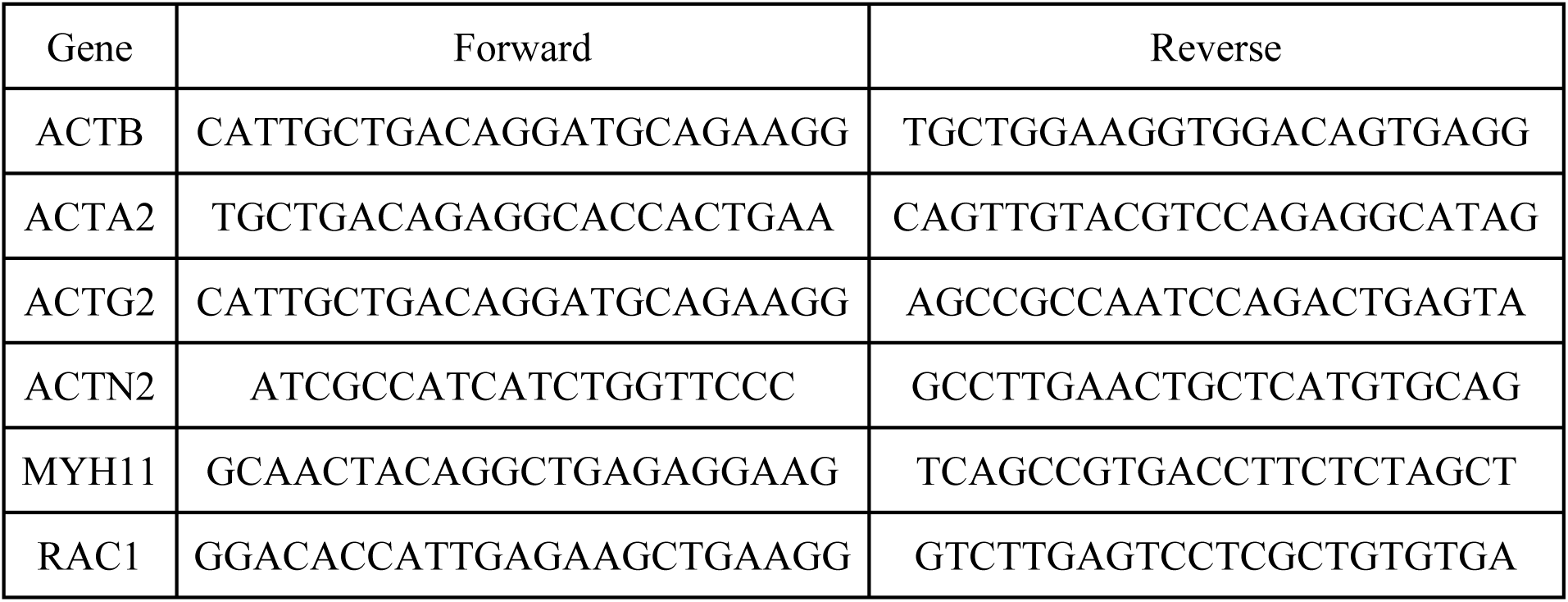
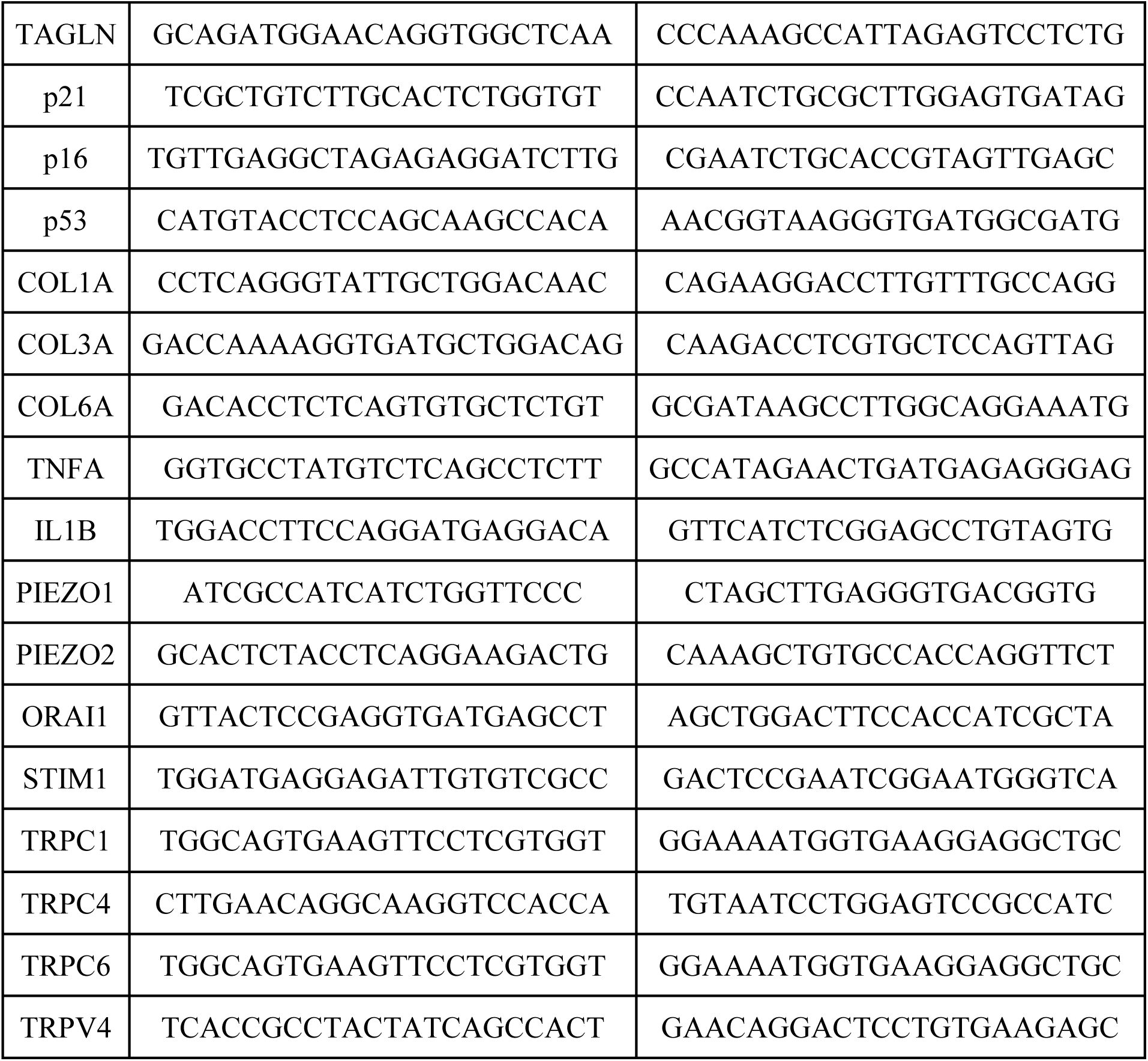

### DNA Microarray analysis

RNA was isolated using RNeasy Mini Kit (Qiagen). Isolated RNA sample quality was assessed by High Sensitivity RNA Tapestation and quantified by Qubit 2.0 RNA HS assay. Reverse transcription was performed based on manufacturer’s recommendation for Affymetrix WT PLUS Kit. The resulting double-stranded cDNA is then amplified and labeled using a Biotin Allonamide Triphosphate. The labeled cDNA was then cleaned up, fragmented, and hybridized to arrays for 16 hr at 45°C. After hybridization, the array was washed on the Fluidics Station 450/250 following the instructions. The probe array is scanned using the GeneChip® Scanner 3000 after the wash protocols were complete. The raw data obtained from the Affymetrix GeneChip Scanner in CEL format was analyzed using Transcriptome Analysis Console (TAC) software that enabled normalization of signal intensity, quality control and statistical analysis of differential expressed genes. The gene lists from the corresponding probe IDs obtained from the TAC software was then used for GO Enrichment Analysis and Kyoto Encyclopedia of Genes and Genomes (KEGG) mapping using the R package Bioconductor (open source) to investigate altered biological pathways associated with aging.

### Calcium imaging

VSMCs seeded coverslips were washed twice using Tyrode solution (1176005, Electron Microscopy Sciences). The cells were then transfected with calcium indicator Fluo-4 (F14201, Thermo Fisher Scientific) for 30 minutes at room temperature, followed by washing with Tyrode solution for 3 times. Imaging of calcium influx were captured using an inverted microscope (Zeiss Axio Observer Z1) at 480 nm excitation wavelength. Images were taken for 30 minutes at 10-s interval. Yoda1 (#21904, Cayman Chemical) at 0.3 uM concentration was added at 2 minutes after recording.

### Image recognition-based quantification of partial deviation

The fluorescent images of F-actin were first skeletonized using a Canny edge detection (MATLAB 2020b). The actin CSK was detected by applying Hough transform (a line extraction technique for digital image processing [71]). Next, the Hough-Peaks associated with the Hough transform matrix were found. Finally, the lines associated with the Hough-Peaks were determined. The angle associated with the lines were used to calculate the orientation of the actin fiber associated with the detected line. The partial actin deviation with the detected lines were then calculated based on formular established in a previous study [45]. Detailed pipeline is presented in **Fig. S8**.

### siRNA transfection

Knockdown of Piezo1 was performed by siRNA transfection with 20 nM of either ON-TARGETplus Piezo1 siRNA Reagents (L-061455-00-0005, Horizon Discovery) or ON-TARGETplus non-targeting control pool (D-001810-10-05, Horizon Discovery) using Lipofectamine™ 3000 Transfection Reagent (L3000001, Invitrogen). VSMCs were cultured in a six-well culture plate at density of 50,000 cells per well overnight, following by addition of Lipofectamine™ 3000 with siRNA reagents according to the manufacturer’s instruction and incubation for 48 hours. The efficiency of Piezo1 knockdown was validated by immunoflorescence and quantitative real-time PCR.

### Single-cell RNA sequencing

Four groups of VSMCs cultured in different conditions (Young, Old, Young + Angiotensin II, Old + siPiezo1) were harvested by enzymatic dissociation and cell viability was confirmed using an automated cell counter (Biorad, TC20). Cell hashing with anti-mouse oligo-tagged antibodies (TotalSeq A, Biolegend) were performed according to the manufacturer’s protocol before mixing them in single-cell suspension. The cell suspension was loaded on a 10x Genomics Chromium instrument to collect individual gel beads in emulsion. The libraries were prepared using Chromium Single Cell 3′ Library & Gel Bead Kit v3.1, PN-1000268, the Chromium Next GEM Chip G Single Cell Kit PN-1000120 and the Dual Index Kit TT, Set A PN-1000215, (10x Genomics). Amplified cDNA was evaluated on a Agilent BioAnalyzer 2100 using a High Sensitivity DNA Kit (Agilent Technologies) and final libraries on a Agilent TapeStation 4200 using High Sensitivity D1000 ScreenTape (Agilent Technologies). Individual libraries were diluted to 2nM and pooled for sequencing. Pools were sequenced with 100 cycle run kits (28bp Read1, 8bp Index1 and 91bp Read2) on the NovaSeq 6000 Sequencing System (Illumina). Cell Ranger Single Cell Software was used to perform de-multiplexing, barcode and UMI processing, and single-cell 3′ gene counting, Seurat package was used for quality control, data filtering, dimensionality reduction, differential gene expression analysis, and uniform manifold approximation.

### Statistics and reproducibility

All data were from at least three independent experiments. Data were first analyzed for normality and then compared with two-sided unpaired t-test using Prism 8 software (GraphPad software Inc.). P-value smaller than 0.05 was considered statistically significant and sample sizes and exact P values are indicated in all figure captions.

## Acknowledgements

We acknowledge financial support from the National Institute of Health (R35GM133646) and the Cancer Center Support Grant (P30CA016087).

## Author Contributions

Ngoc Luu, Apratim Bajpai, Conceptualization, Data curation, Formal analysis, Visualization, Investigation, Methodology, Writing – original draft; Rui Li, Seojin Park, Mahad Noor, Data curation, Formal analysis, Writing – review and editing; Xiao Ma, Methodology, Writing – review and editing; Weiqiang Chen, Conceptualization, Formal analysis, Supervision, Funding acquisition, Investigation, Methodology, Writing – original draft, Writing – review and editing, Project administration;

