## Supplementary material for "Aging-associated Decline in Vascular Smooth Muscle Cell Mechanosensation is Mediated by Piezo1 Channel": SI Material

**This Supplementary Information file includes:**

Supplementary Figures S1-S11.

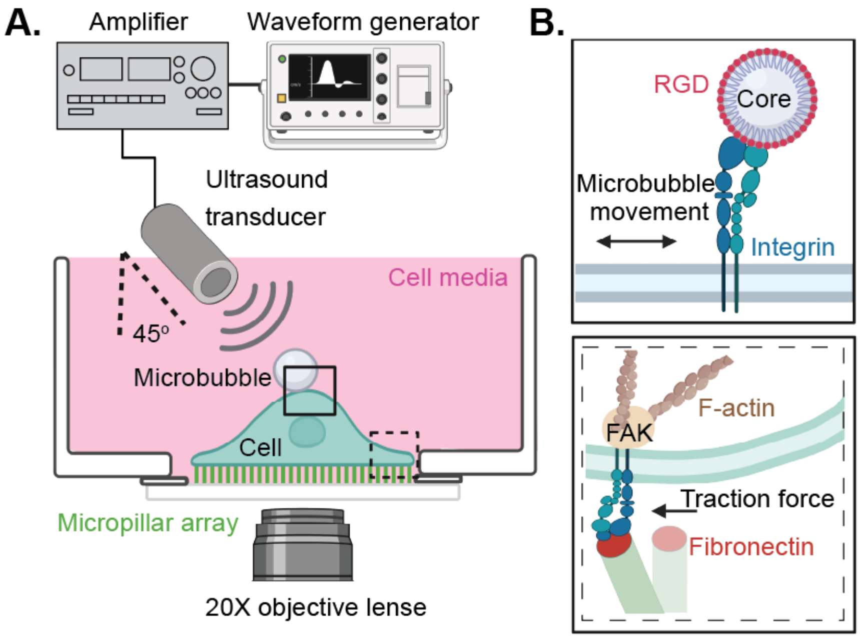


**Figure S1. Schematic illustration of the ultrasound tweezers-based micromechanical system for single-cell mechanosensation study. A)** Experimental apparatus including a function generator, a power amplifier, and an ultrasound transducer that applies ultrasound pulses to generate acoustic radiation force on the microbubble attaching on cell membrane causing its displacement. The cell is loaded on a PDMS micropillar array substrate and the cellular force response is observed with an inverted microscope. **B**) Schematic illustrations showing attachment of microbubble onto cell membrane via RGD-integrin binding (top panel), and cell traction force measurement based on the deflections of micropillars underneath the cell (bottom panel). Figure is created with BioRender.com.

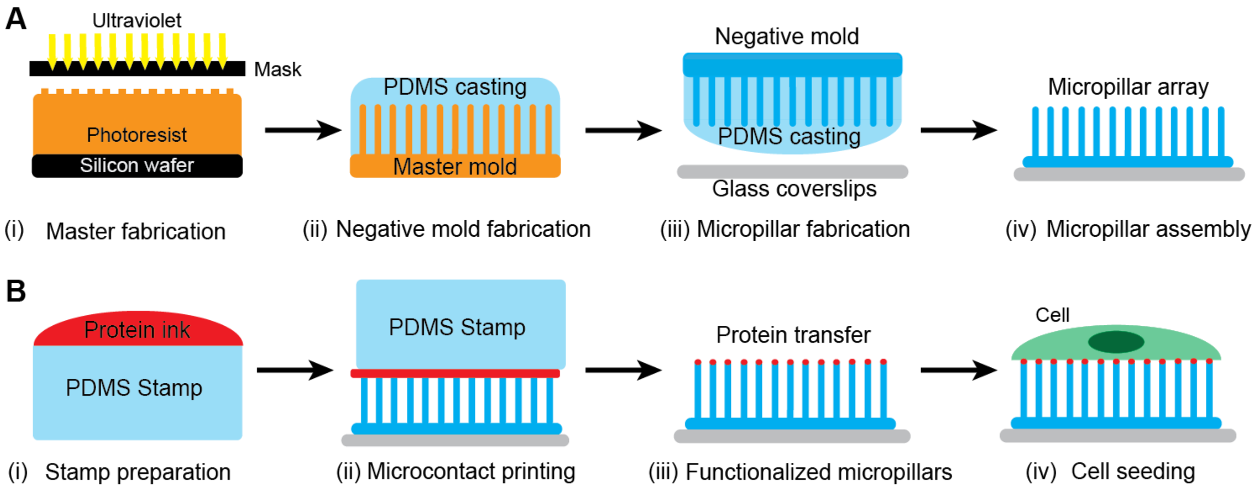


**Figure S2: Micropillar array fabrication, microcontact printing, and cell seeding**. **A**) Schematic illustration of the PDMS micropillar array fabrication procedure which includes (i) silicon master mold manufacturing using photolithography, (ii) creating PDMS negative mold, and (iii-iv) making PDMS micropillar array on a glass substrate from the negative molds though a soft lithography fabrication process. **B**) Schematic illustration of the cell seeding process, including (i) coating the PDMS stamp with protein solution containing fibronectin and fluorescence-conjugated fibrinogen adhesive proteins, (ii-iii) transferring the adhesive proteins onto the tips of the micropillars using a microcontact printing technique, and (iv) loading of cells onto the functionalized micropillar array substrate.

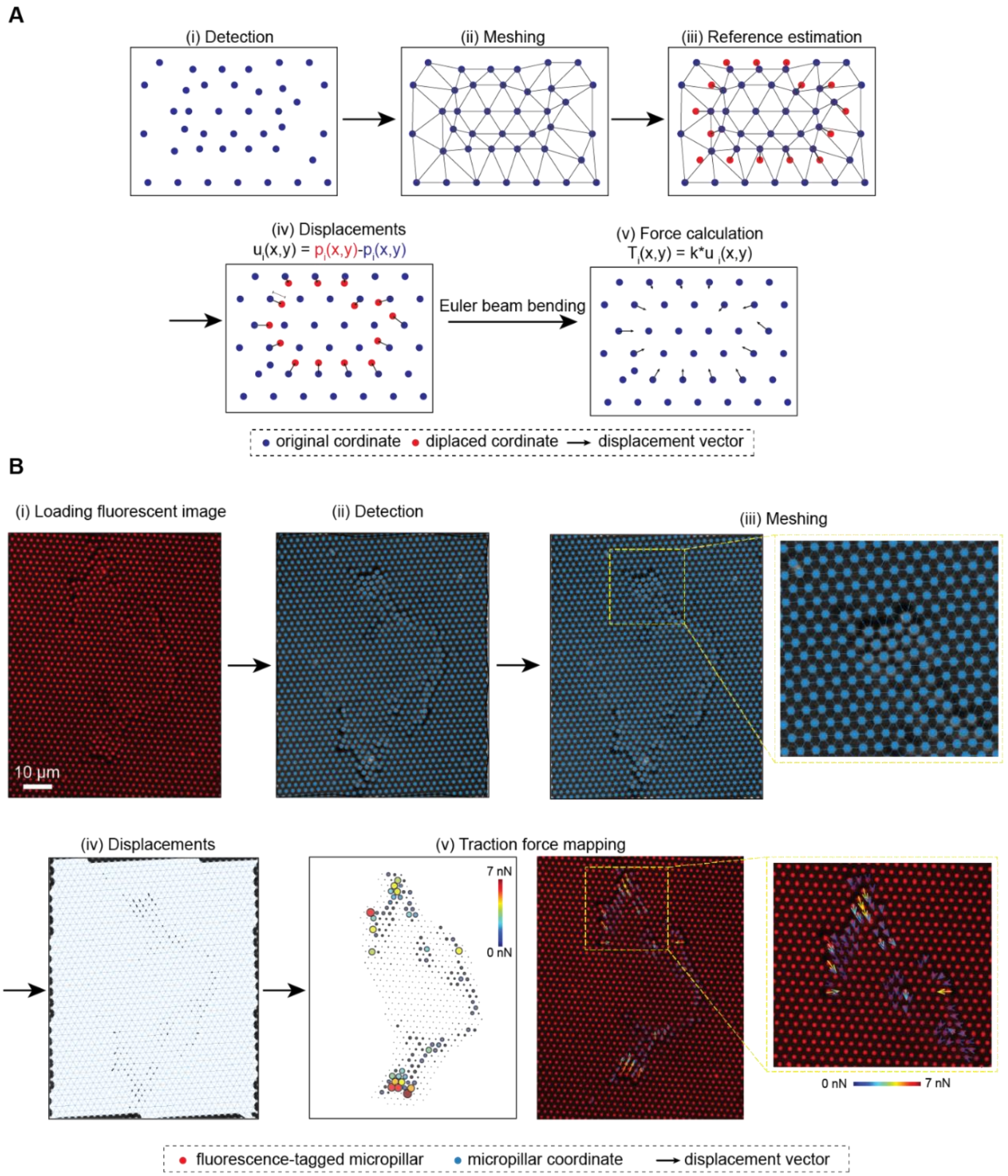


**Figure S3: Traction force measurement using the PDMS micropillar array. A**) The pipeline stages showing calculation of traction force using Cellogram software (open source). The software first detects the coordinate of single micropillar and presents them as a circle dot (i). Next, a mesh is generated to connect all the detected micropillars (ii). Energy minimization is performed to detect the reference position of the micropillars (iii). Displacement vector is obtained by subtracting the displaced coordinates with the reference coordinates. Finally, an elastic beam theory based on displacement and spring constant of the micropillars is applied to calculate the traction forces exerted by the cells. **B**) Example of the workflow showing traction force mapping from the fluorescent images (red channel) of the micropillars. The v panel shows a representative traction force heatmap (left) and a traction force vector map (right) of VSMC.

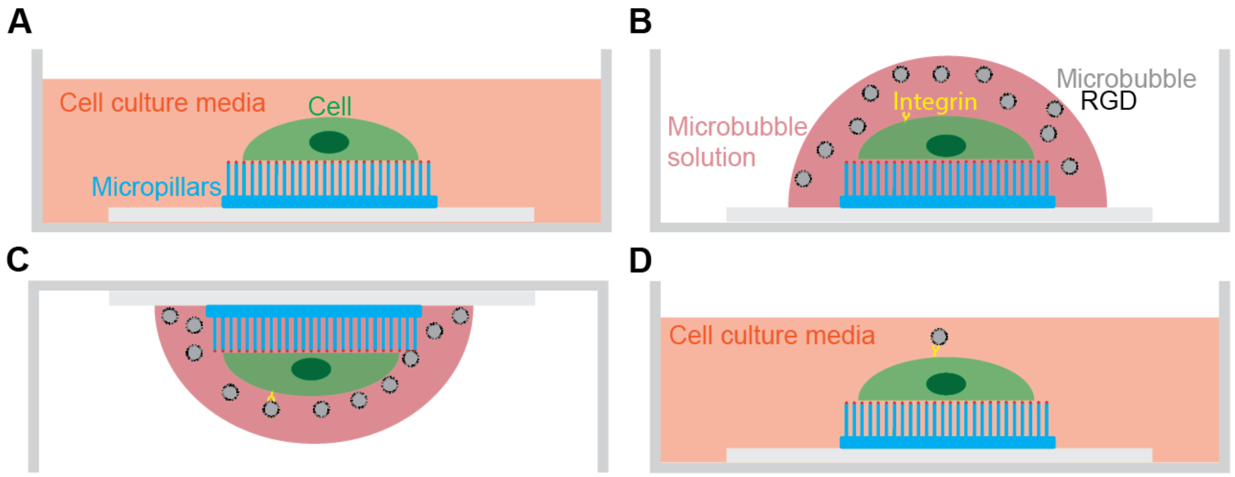


**Figure S4: Microbubble attachment to cell**. **A**) Single cells are seeded on a fibronectin-coated micropillar array substrate. **B**) Cell culture media is removed, and microbubble solution was added to the cells. **C**) The culture dish is flipped for 20 minutes to facilitate microbubble attachment via floatation. **D**) Unbounded bubbles are washed away with cell culture media twice, and cells bound with microbubbles are then suspended in fresh culture media for following ultrasound stimulation experimentation.

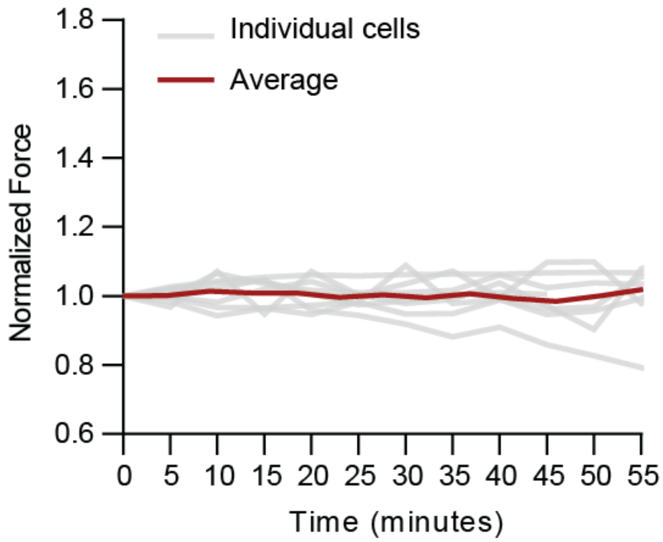


**Figure S5**: **Temporal evolution of normalized traction force of VSMCs on PDMS micropillar array without ultrasound stimulation.** Gray lines represent temporal force response of individual cells, and red line represents average force response of all measured cells (n = 8).

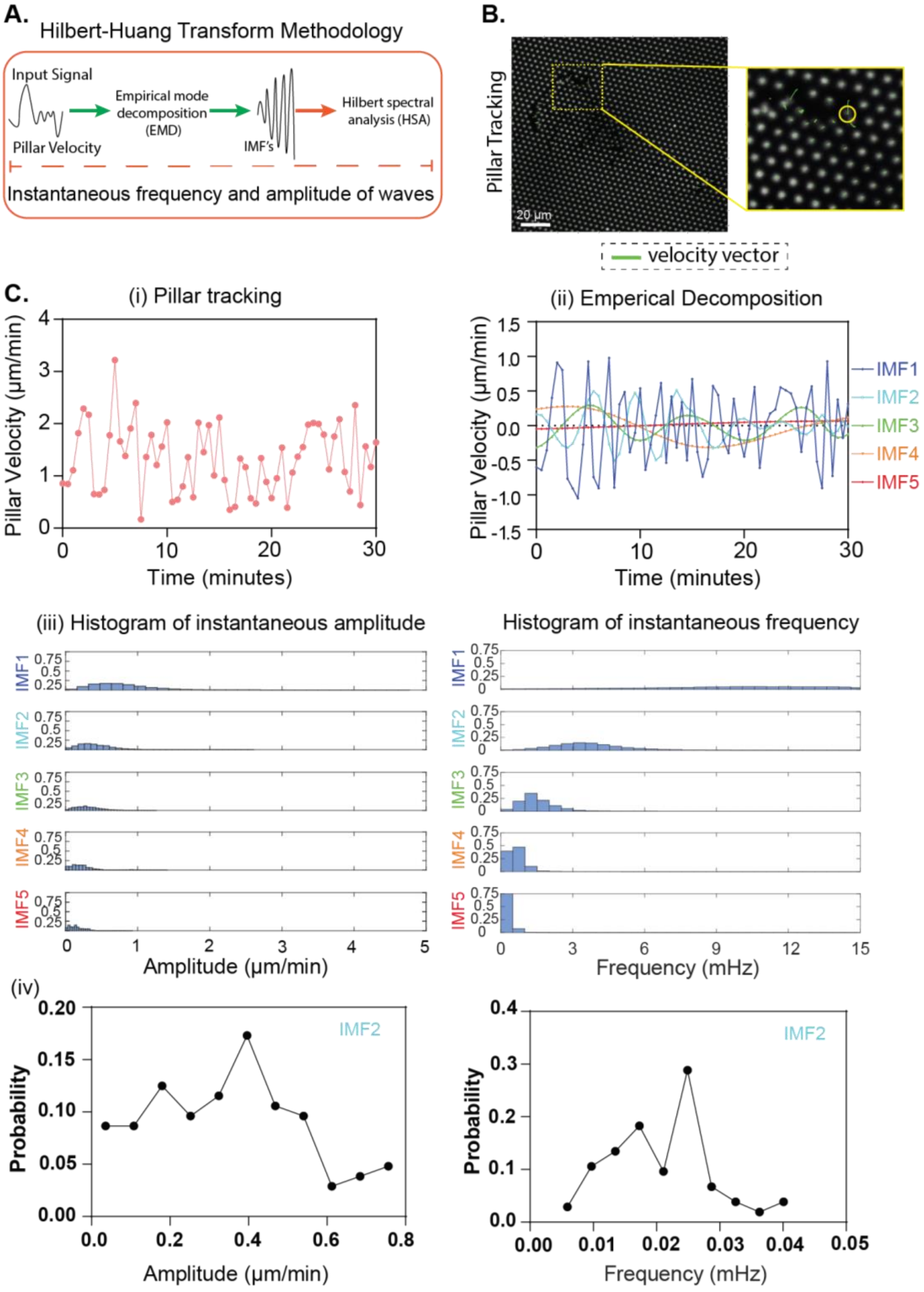


**Figure S6: Instantaneous frequency spectrum analysis. A)** Pipeline of Hilbert-Huang transform methodology, including tracking individual micropillar's velocity, performing empirical mode decomposition (EMD) of the velocity signal, and then applying Hilbert transform to the decomposed signal to get the instantaneous spectrum**. B)** Representative images showing displacement of individual pillars (white dots) and velocity (green arrows)**. C)** Application of EMD on the pillar velocity**. (i)** Pillar velocity variation over time serving as the input signal**. (ii)** EMD separates the velocity time series into a finite and small numbers of component signals called intrinsic mode functions (IMFs). The number of IMFs was predetermined as 5 in this study based on the termination criterion, i.e. the variance of residual signal (equals to subtraction of the sum of IMF1 to IMF5 from the original signal) is less than 5% of that of the original signal. Hilbert Transform is applied to each IMF to compute instantaneous amplitude and instantaneous frequency of the micropillar's velocity signal. **(iii)** Histograms showing the instantaneous amplitude and instantaneous frequency in the signal from different IMFs. (iv) Examples of probability distributions of instantaneous properties (amplitude and frequency) derived from IMF2 of the micropillar's velocity signal.

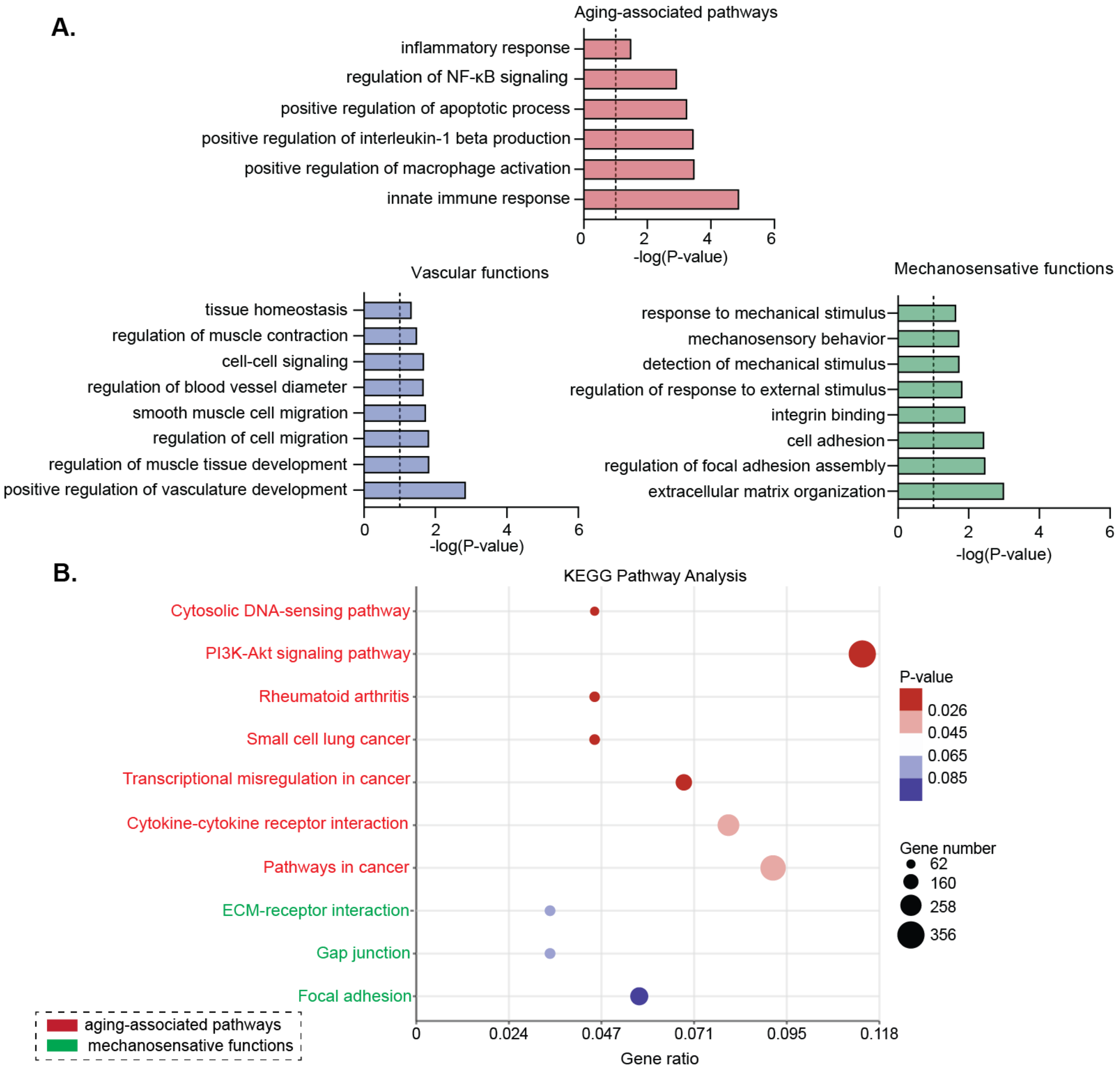


**Figure S7. Pathway analysis of differentially expressed genes in young versus old VSMCs. A)** GO enrichment analysis demonstrating alterations in VSMC biological processes with aging, based on the Gene Ontology Resource knowledgebase. The enriched pathways were categorized into aging-associated pathways, vascular functions, and mechanosensitive functions. **B)** KEGG pathway analysis performing on differentially expressed genes in young and old VSMCs (n = 3 for each group).

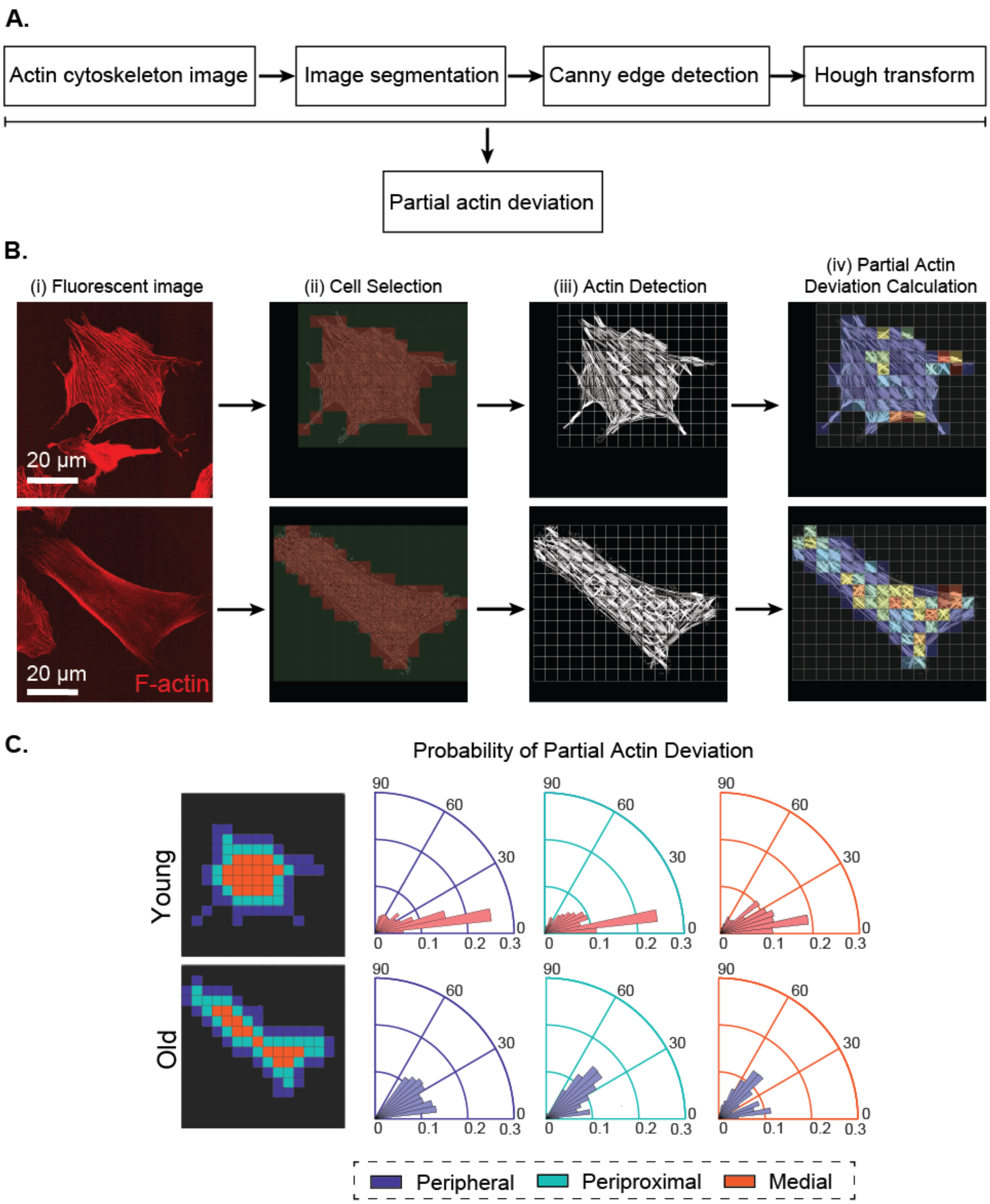


**Figure S8. Subcellular F-actin analysis. A)** Outline of image recognition-based actin CSK quantification method to calculate partial actin deviation. **B**) An example demonstrating subcellular actin analysis of young and old cells, including raw F-actin fluorescent image (i), cellular selection using boxes of 128x128 pixel (ii), F-actin fiber detection (iii), and partial actin deviation calculation. **C**) Polar histogram quantifying the subcellular distribution of the PADs in peripheral, periproximal and medial section of young and old cells as indicated.

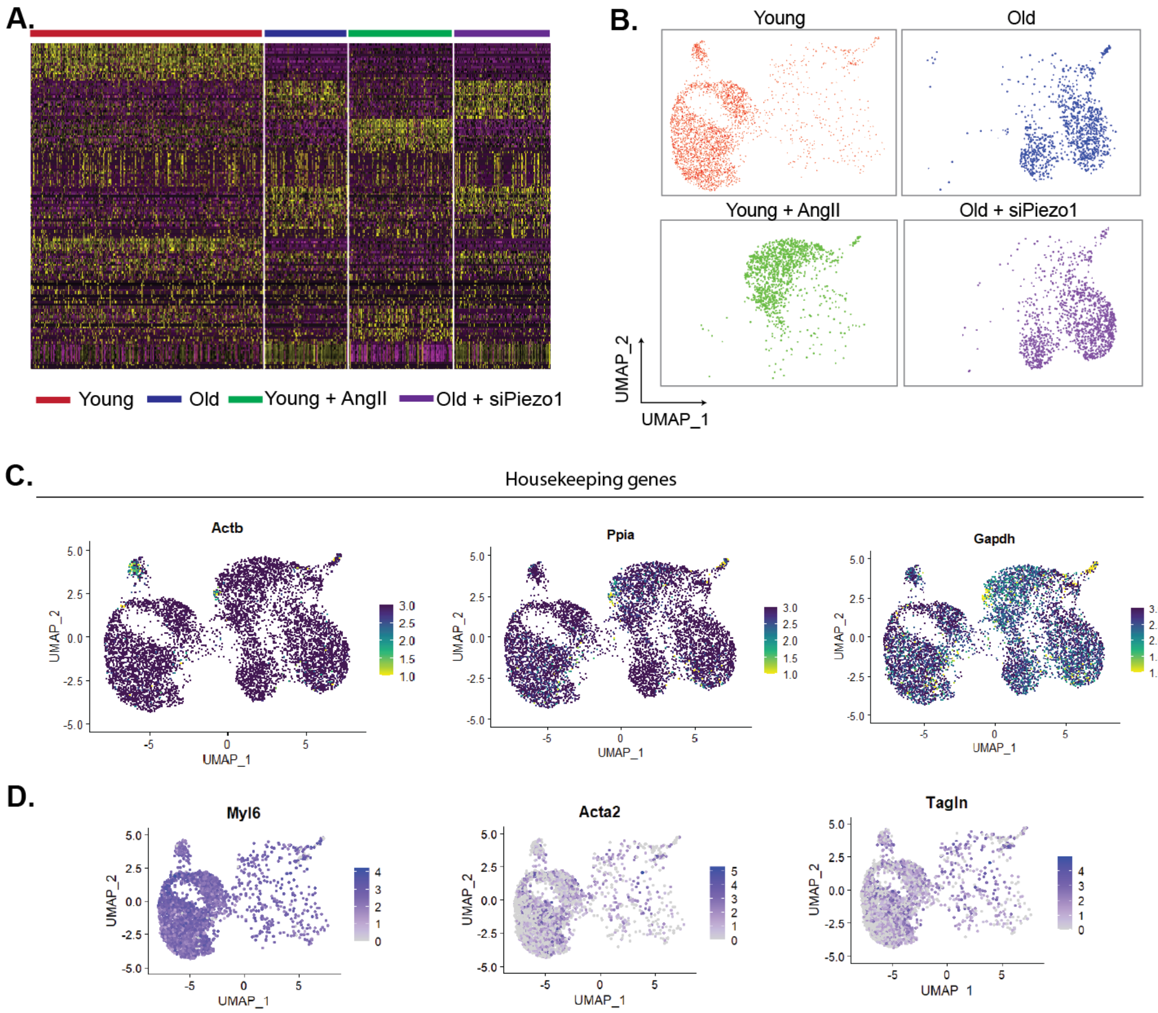


**Figure S9. A)** Heatmap representation of top 20 differentially expressed genes across four cell clusters**. B)** UMAP projection of each VSMC subset labeled by obligo-tagged antibodies**. C)** Feature plots showing expression of housekeeping genes across the VSMC subsets. **D**) UMAP projection highlighting the expression of VSMC markers in cluster of young cells.

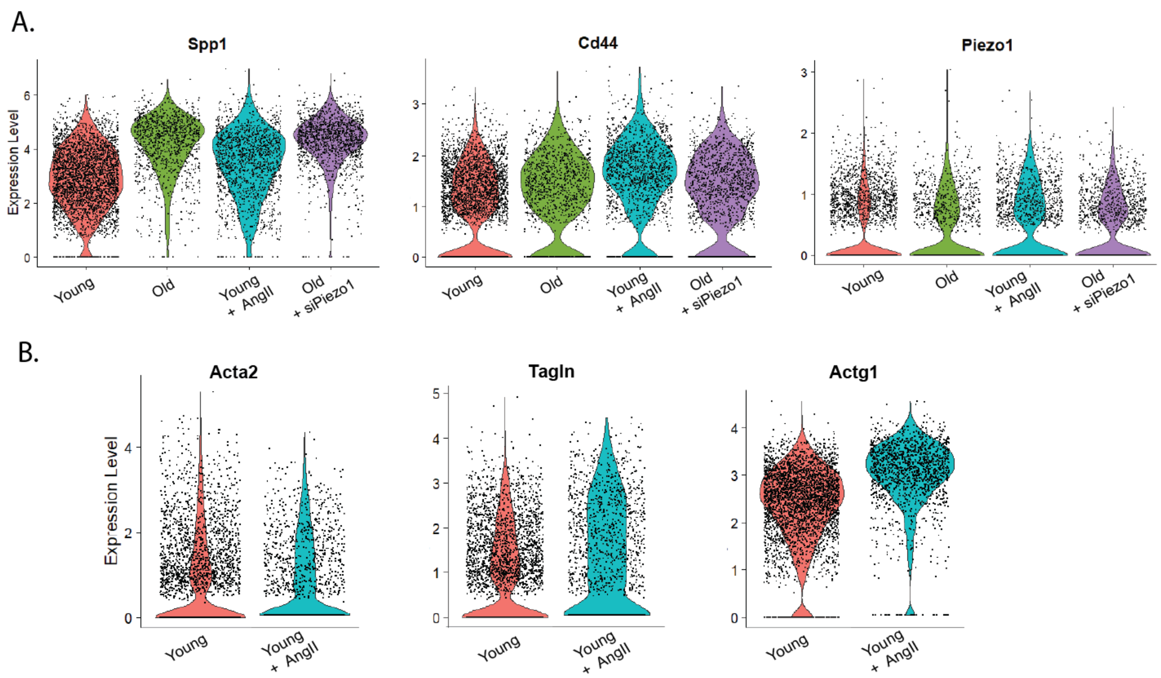


**Figure S10. A)** Violin plots for markers of Spp1, Cd44, and Piezo1 among the four clusters. **B**) Violin plots quantifying expression of cytoskeletal genes in young versus young AngII cluster.

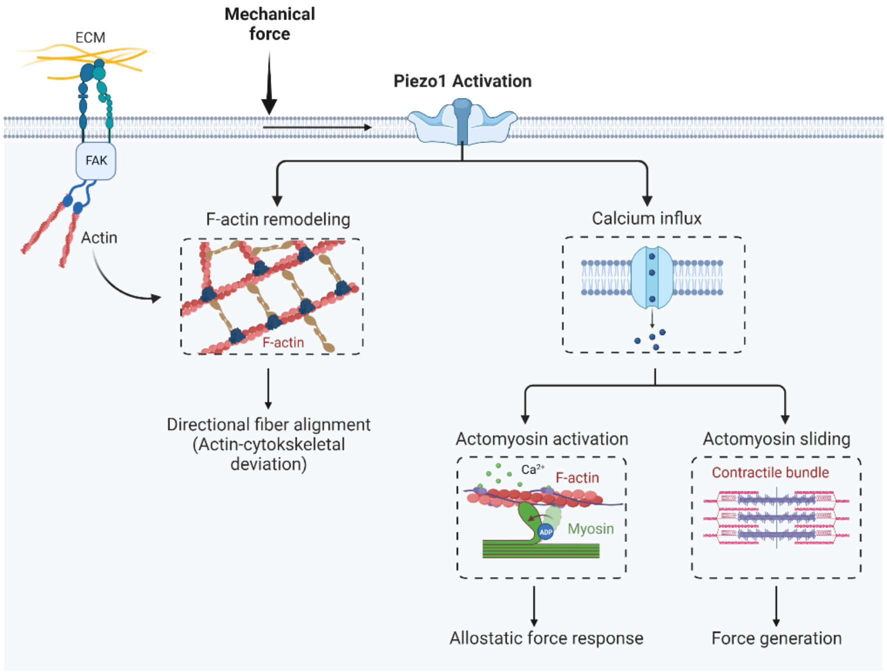


**Figure S11. Schematic of Piezo1-dependent signaling pathway that mediates VSMC mechanosensation.** Figure is created with BioRender.com.
